# Integrating Epstein-Barr virus (EBV) status into diffuse large B cell lymphoma (DLBCL) genetics

**DOI:** 10.64898/2026.04.03.710620

**Authors:** Quincy Rosemarie, Mitchell Hayes, Eric C Johannsen

## Abstract

Diffuse large B-cell lymphoma (DLBCL), the most common aggressive lymphoma, encompasses histologically similar but genetically distinct cancers. Recent genetic studies have defined at least six molecular subtypes, yet none account for Epstein-Barr virus (EBV), despite 5-15% of DLBCLs being EBV-associated. By reanalyzing published whole-exome and RNA-sequencing data from 481 tumors, we identified 19 EBV-positive cases. These were significantly enriched in the BN2 subtype (6/19), while most (11/19) remained unclassified. In BN2 tumors, several subtype-defining mutations were reduced in frequency among EBV-positive cases, supporting the hypothesis that EBV oncogenes substitute for specific cellular alterations and may confound DLBCL classification algorithms. Extending our analysis to cell lines, we found that the widely used Val cell line harbors the B95-8 laboratory EBV strain; other EBV-positive lines appeared authentic but modeled only non-BN2 subtypes and expressed an atypical viral latency III program, whereas some DLBCL tumors expressed the atypical latency III program and others latency I or II. Together, these findings demonstrate that EBV-positive DLBCL, like DLBCL itself, is not a single disease, and that current *in vitro* models only partially capture its biological heterogeneity.

**Key points:**

- EBV-positive DLBCL is not a single disease and EBV status can impact genetic-based classifications.
- Current EBV-positive DLBCL cell lines do not adequately capture tumor complexity; we determined that Val is a problematic cell line.

## Introduction

Diffuse large B cell lymphoma (DLBCL) is the most common aggressive lymphoma, and 5-15% of cases are attributable to Epstein-Barr virus (EBV) infection. DLBCL is not a single disease, but a group of histologically similar yet genetically distinct cancers. DLBCL nosology has improved both prognostication and therapy. For example, the cell-of-origin (COO) classification, developed using gene-expression profiling, distinguishes activated B-cell–like (ABC) DLBCL – which carries a poorer prognosis – from germinal center B-cell-like (GCB) DLBCL^1,2^. More recently, genetic classifications based on recurrent patterns of cellular mutations have largely converged on ∼6 subtypes^3–6^. None of these, including the widely adopted LymphGen classifier, consider EBV. This omission limits our understanding of which DLBCL subtypes are causally associated with EBV and, because EBV is a dependency factor for its associated lymphomas, has implications for therapy.

Distinct from these efforts are studies characterizing the genomic and transcriptomic landscapes of EBV-positive DLBCL. Several studies have reported elevated JAK/STAT and NF-κB signatures relative to EBV-negative controls^7–9^. Targeted and unbiased sequencing of EBV-positive DLBCL tumors have identified enriched mutations relative to EBV-negative DLBCL, although only a few genes were frequently mutated across more than one study. These include Notch2, KMT2D, ARID1A, STAT3, SOCS1, and TET2. This inconsistency is attributable in part to the implicit assumption that EBV-negative DLBCL represents a single disease with consistent baseline gene mutation frequencies. As a result, changes in mutation frequency attributed to EBV may instead reflect differences in the prevalence of subtypes within the EBV-negative cohort. Furthermore, EBV-positive DLBCL itself may not represent a single disease. Few studies have attempted to determine whether EBV-positive DLBCL corresponds to any of the genetic subtypes defined for EBV-negative DLBCL^10,11^. These limited analyses reveal that the majority of EBV-positive DLBCL remain unclassified by algorithms such as LymphGen, leaving open the question whether these “unclassified” tumors are related to each other.

The patterns of EBV latent gene expression in EBV-positive DLBCL provide additional evidence that it is not a single disease entity. Most EBV associated cancers exhibit characteristic patterns of EBV latent gene expression. Latency I, in which only one protein coding gene (EBNA1) is expressed, is seen in Burkitt lymphoma (BL) and EBV-associated gastric carcinoma (EBVaGC)^12^ EBV associated Hodgkin lymphoma and nasopharyngeal carcinoma exhibit latency II, in which LMP1, LMP2A, and LMP2B are expressed in addition to EBNA1^13^. The full repertoire of EBV latent genes (latency III) is seen most commonly in immunodeficiency-associated cancers such as post-transplant lymphoproliferative disease or AIDS-associated primary CNS lymphoma^14,15^. By contrast, EBV-positive DLBCL lacks a typical latency pattern: most studies observe latency II or latency III, but a significant minority of cases express the latency I program^16–19^. EBV gene expression may also complicate efforts to classify DLBCL genetically. For example, studies of EBV-positive nasopharyngeal carcinoma (NPC) have found that LMP1 expression substitutes for NF-kB mutations to such an extent that they are mutually exclusive^20^. Thus, EBV-positive DLBCL may lack some of the driver mutations on which DLBCL classification systems depend. It is therefore essential to define which EBV oncogenes are expressed and how they influence somatic mutation profiles in DLBCL.

We investigated the genetic landscape of EBV-positive DLBCL by reanalyzing published DLBCL RNA-seq data to assess whether specific subtypes are associated with EBV and to define patterns of EBV latent gene expression and their interactions with recurrent host mutations. We complement this analysis by examining the current repertoire of EBV-positive DLBCL cell lines to determine whether they faithfully model EBV gene expression patterns and host mutations observed in EBV-positive tumors. Our results provide a framework for understanding how *in vitro* models relate to actual tumors and highlight the need to establish additional cell lines to represent the diversity of this EBV-associated disease.

## Results

### Identification of EBV-positive DLBCL tumors

We reanalyzed RNA sequencing data from 481 DLBCL tumors^3^ and quantified reads mapping to the EBV genome. Most tumors had fewer than 1 EBV read per million mapped, and we tentatively identified 20 candidate EBV-positive DLBCL based on EBV read counts > 5 per million mapped (Figure 1A). One tumor exhibited only lytic gene expression and was excluded from further analysis, as the absence of EBV latent gene expression is inconsistent with an EBV-positive tumor (Figure 1B, Supplementary Figure S1). We classified the 19 tumors into one of three canonical latency programs: 2 latency I (11%), 9 latency II (47%), and 8 latency III (42%). This heterogenous latent gene expression of DLBCL contrasts the stereotyped latency gene patterns seen in most other EBV-associated cancers^21–26^.

**Figure 1.**
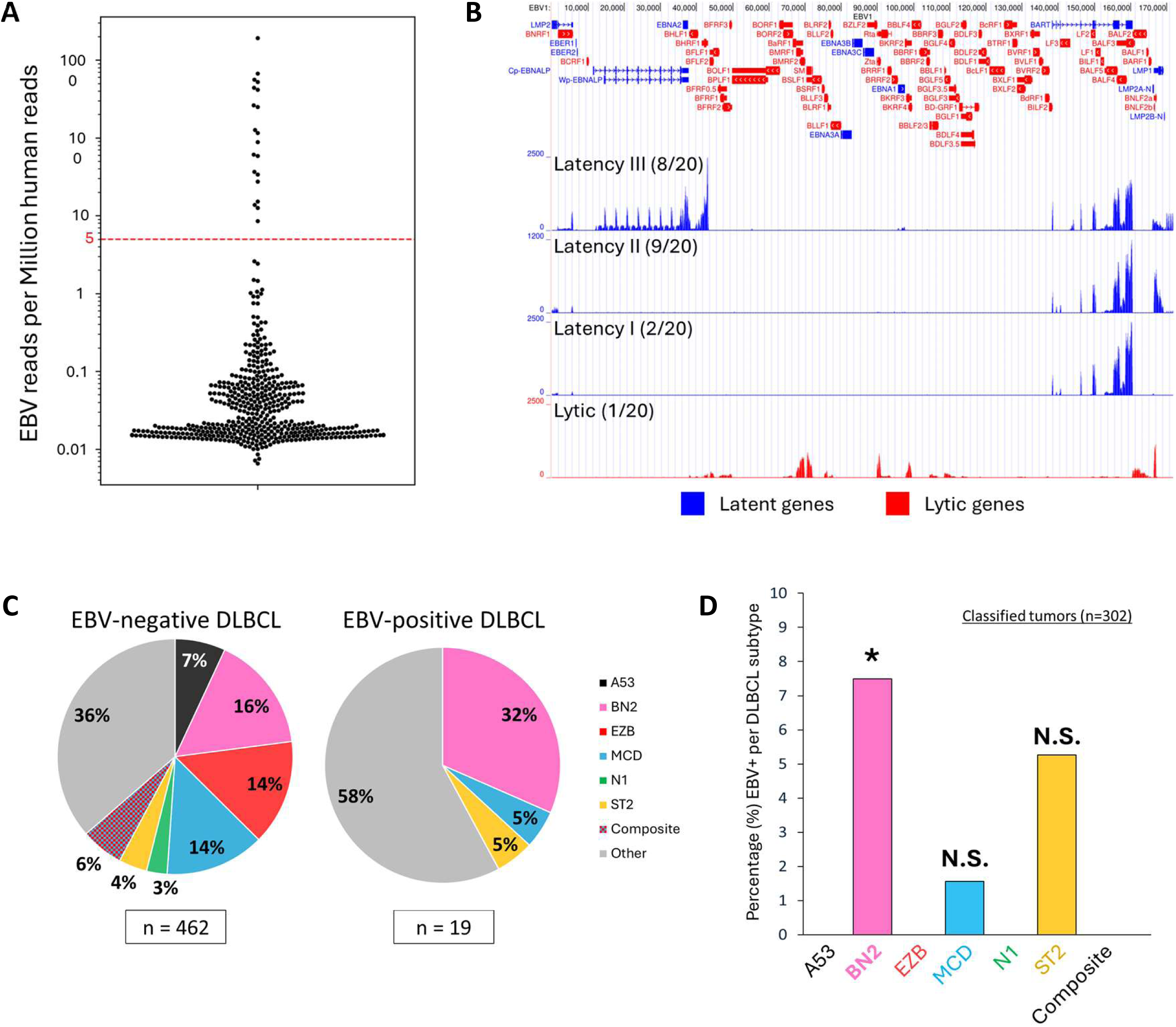
EBV-positive DLBCLs have a distinct genetic subtype distribution. A) Dot plot showing EBV reads per million human reads from DLBCL tumors (n=481). Initially, we chose a threshold of 5 reads/million as a cut-off, resulting in 20 candidate EBV-positive DLBCLs. B) Representative tracks from candidate EBV-positive DLBCL tumors displayed on the UCSC genome browser. Genes expressed during latency (blue) or lytic replication (red) are indicated. Examples of EBV-positive tumors expressing each latency program (I, II, or III) are shown. One tumor biopsy had reads mapping only to lytic genes and was excluded from further analysis. C) Proportions of LymphGen subtypes in EBV-negative and EBV-positive DLBCL tumors. BN2 is a prominent subtype among the EBV-positive DLBCL and most of the remaining tumors are unclassified (‘Other’). D) Bar chart showing percentage of EBV-positive tumors for each subtype. Among the 302 tumors classifiable by LymphGen, EBV-positive DLBCL is enriched within the BN2 subtype (*: p-val < 0.05, 2×2 Barnard’s exact test with F-order; each subtype is compared to the rest of the classified subtypes, e.g., BN2 vs non-BN2).

### EBV-positive DLBCL is enriched within the BN2 subtype and many are unclassified

Most EBV-positive DLBCLs were BN2 subtype (n=6) or unclassified (Other, n=11) by the LymphGen algorithm (Figure 1C, Table S1). Additionally, one EBV-positive tumor was MCD subtype and another was ST2. Among the classifiable tumors (i.e., excluding Other), only BN2 showed a statistically significant enrichment of EBV-positive tumors (Figure 1D). This finding indicates that a subset of EBV-positive DLBCL has a shared mutation profile, and possibly a shared pathogenesis, with BN2 DLBCL. The unclassified (Other) EBV-positive DLBCL may constitute a novel genetic subtype.

Most EBV-positive cancers have stereotyped latent gene expression programs. For example, nasopharyngeal carcinoma and Hodgkin lymphoma express the latency II program, whereas BL typically express the highly restricted latency I program (and less frequently Wp- restricted)^23^. We found that BN2 subtype tumors expressed latency II or III (Supplementary Table S1). The Other subtype, included tumors with latency I, II, and III gene expression. The EBV-positive MCD tumor and the ST2 tumor each expressed Latency III. Thus, even within specific subtypes of DLBCL, EBV latent gene expression appears to be heterogenous.

### EBV may confer a worse prognosis in specific DLBCL subtypes

EBV-positive tumors are associated with a poor prognosis in some but not all studies^7,27–31^. We therefore examined how EBV status affects outcomes in this cohort. When EBV status was considered regardless of DLBCL subtype, there was a trend towards worse OS and PFS, that did not reach statistical significance (Supplementary Figure S2). When DLBCL subtype was also considered, we found EBV conferred a significantly worse prognosis for BN2, but not for the ‘Other’ subtype. While these results are preliminary, they indicate that the impact of EBV on prognosis may be subtype dependent.

### Influence of EBV on DLBCL mutations

EBV-positive lymphomas are dependent on the virus, and do not survive experimental EBV eviction^32–34^. Moreover, because the EBV genome is maintained inefficiently in dividing cells^35^, proliferating cells lose the virus unless it confers a growth or survival advantage. Thus, EBV oncogenes promote lymphomagenesis and likely reduce the number of oncogenic mutations required for DLBCL. Indeed, many viral cancers have a lower mutational burden than their virus-negative counterparts^36,37^. To identify candidate gene mutations that EBV may replace, we compared the mutation profiles of EBV-positive and EBV-negative tumors. As it is unclear whether the ‘Other’ DLBCL tumors represent a single disease, we focused this analysis on the BN2 subtype.

In BN2 DLBCLs, several subtype-defining genes were mutated at reduced frequencies in the EBV-positive compared to the EBV-negative tumors (Figure 2A). The impact of EBV status was not limited to subtype defining genes. Many genes were found to be mutated in EBV-positive DLBCL at frequencies that differed substantially from their EBV-negative counterparts (Figure 2B, Supplementary Figure S3A). Notably, EBV-negative BN2 harbored gain-of-function mutations in BCL10 and CARD11 as well as loss of function in NFKBIE. Each of these mutations leads to aberrant NF-κB signaling, and their absence in EBV-positive BN2 suggests that NF-kB signaling by viral LMP1 and/or LMP2A supplants the need for these oncogenic mutations. In the EBV-positive BN2 tumors, CD70 mutations were observed in all tumors expressing the latency III program, but absent in those expressing the more restricted latency II (Supplementary Figure S3B). CD70 is known to be important for immune responses to EBV infection^38,39^. Loss of CD70 function may provide an alternative mechanism to evade immune responses in tumors that remain dependent upon the full repertoire of EBV oncogenes expressed in latency III.

**Figure 2.**
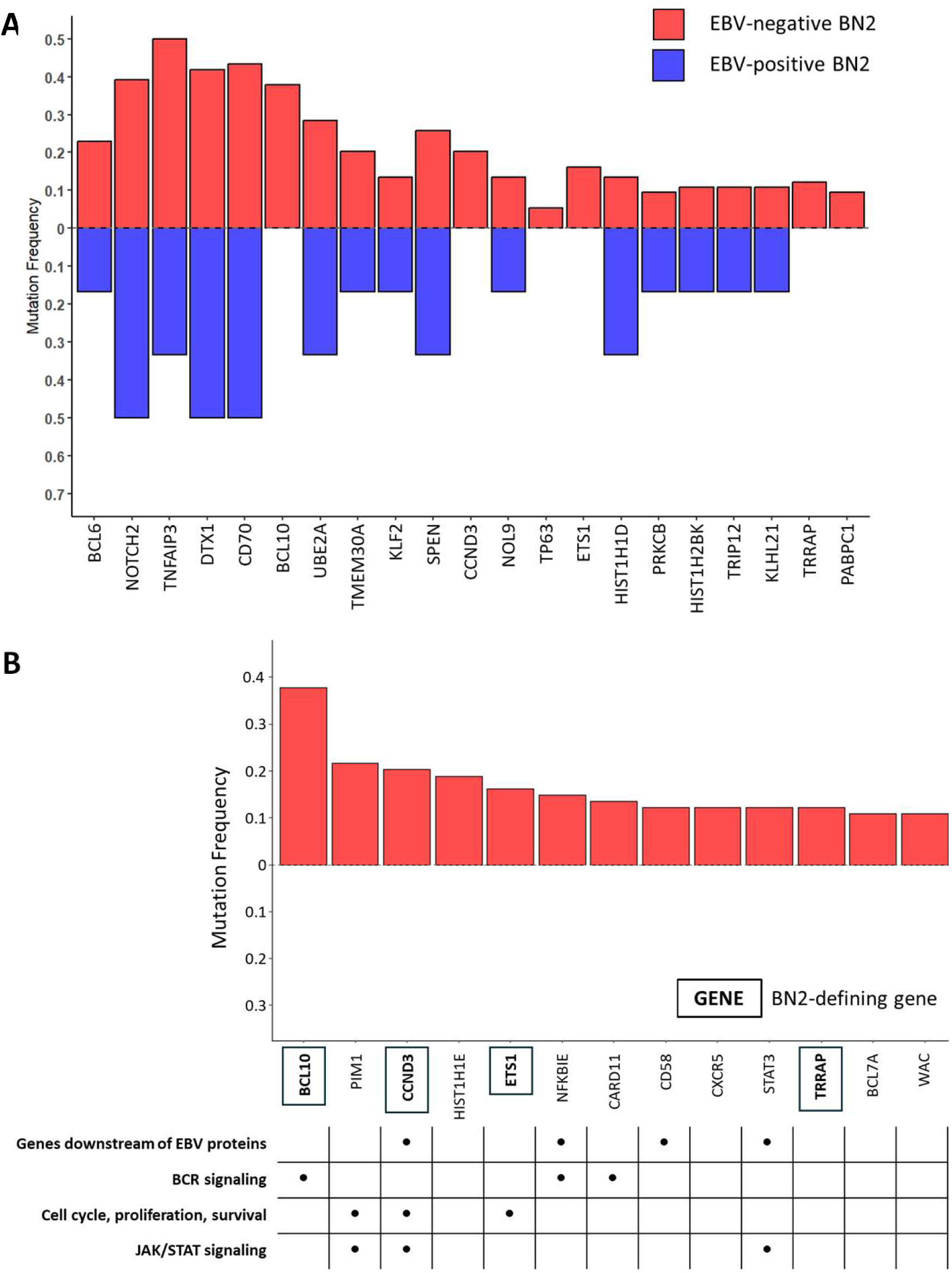
Cellular mutation frequency differs with EBV status. A) Mutation frequencies of BN2-defining genes in EBV-negative and EBV-positive BN2 tumors. B) Genes not mutated in EBV-positive BN2 with a mutation frequency ≥ 0.1 in EBV-negative BN2.

Among the mutated genes that are enriched in EBV-positive BN2 tumors are the epigenetic regulators, ARID1A and KMT2D (Supplementary Figure S3A).^40^. Both genes are frequently mutated in EBV-positive DLBCL^10,41^, and other EBV-positive malignancies.^42–46^.

### RNA-seq is consistent with EBV-positive DLBCL having increased tumor-infiltrating immune cells

Transcriptional profiling underlies COO subtyping, but does not recapitulate genetic-based classification of DLBCL with unsupervised clustering. This likely reflects substantial contributions from infiltrating non-malignant cells that obscure tumor intrinsic transcription differences. Based on their previously reported COO classification,^3,4^ we found most of EBV-positive DLBCL were non-GCB subtype (Figure 3A) as has been found in other studies^7,10,47^. We next used CIBERSORTx^48^ to infer their immune cell composition. Within the BN2 and Other genetic subtypes, EBV-positive cases exhibited increased proportions of non-B, tumor-infiltrating immune cells, mainly consisting of T cells and macrophages (Figure 3B & 3C). Previous reports have observed such infiltrates in EBV-positive DLBCL. Similar immune-rich microenvironments have been described in previous studies of EBV-positive DLBCL^28,49,50^.

**Figure 3.**
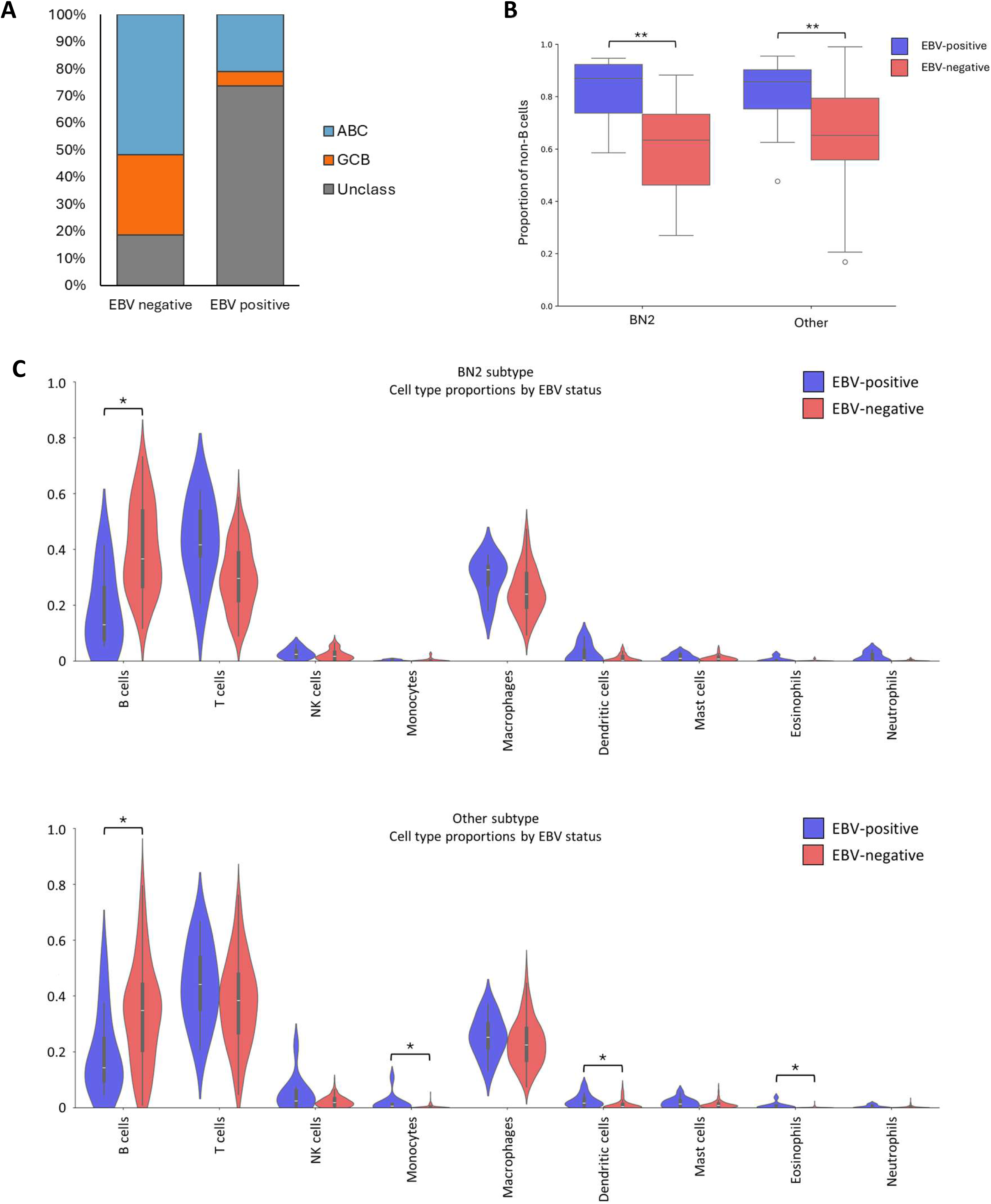
RNA-seq of EBV-positive DLBCL is consistent with elevated immune infiltration. A) Bar chart showing the proportion of cell-of-origin (COO) subtypes in EBV-negative and EBV-positive DLBCL. ABC: activated B cell-like, GCB: germinal center B cell-like, Unclass: unclassified. B) Box plots indicating increased non-B cells in EBV-positive tumor biopsies relative to EBV-negative for both BN2 and Other subtypes. C) Violin plots indicating that tumor-infiltrating immune cells in DLBCL consist mainly of T cells and macrophages (*: p-val < 0.05, **: p-val < 0.01, Mann-Whitney U test).

### Some unclassified EBV-positive DLBCLs approximate defined subtype(s) - others may represent a novel subtype

The EBV-positive DLBCLs have significantly fewer genetic alternations than the EBV-negative tumors (Figure 4A), as has been observed with other EBV-associated malignancies^36,37^. To examine whether the “Other” EBV-positive tumors would be classifiable if EBV oncogenes were accounted for, dimensionality reduced LymphGen features were used to visualize the proximity of EBV-positive tumors to the network of classified tumors (Figure 4B). The result broadly recapitulates the separation of LymphGen subtypes and several “Other” EBV-positive tumors were near the BN2 cluster. Most of the remaining EBV-positive “Other” tumors formed a cluster, potentially representing a novel subtype. Analysis of larger cohorts of EBV-positive tumors is needed to resolve these important issues.

**Figure 4.**
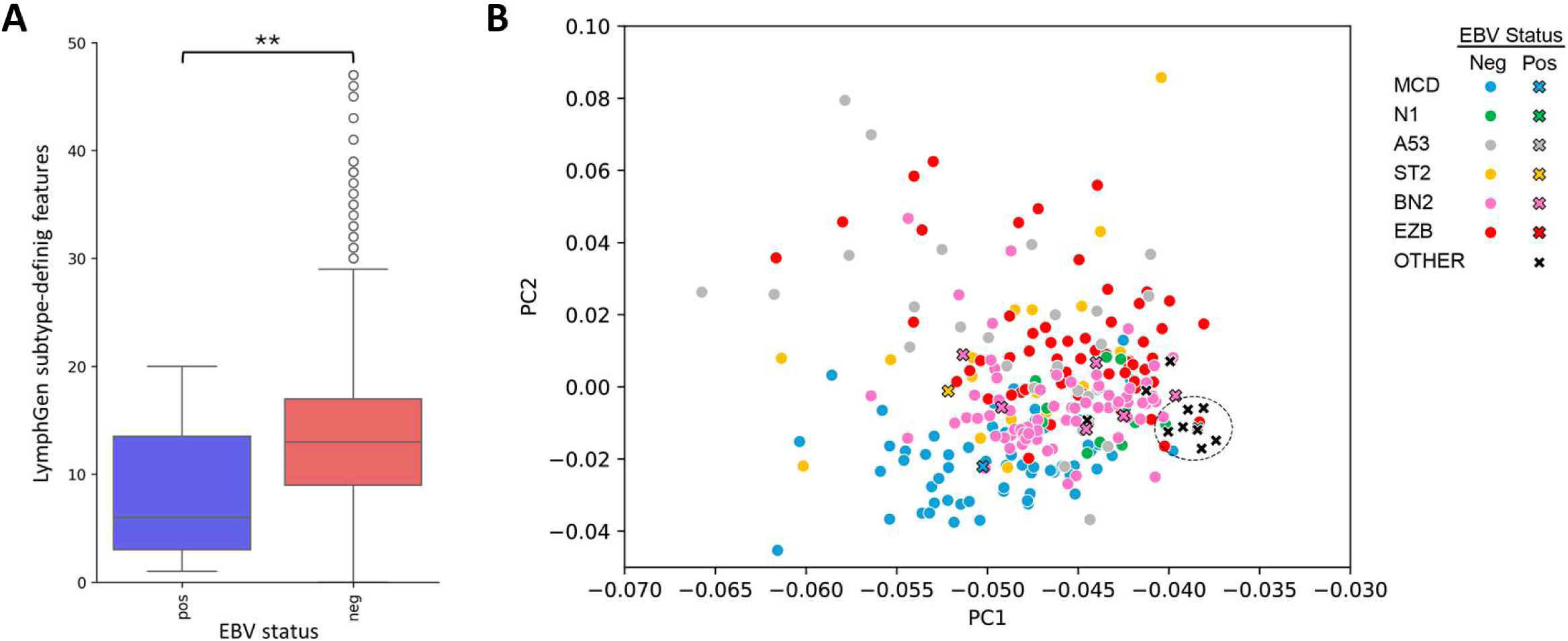
EBV-positive DLBCL form a cluster. A) Bar chart showing the number of LymphGen subtype-defining features in EBV-positive and EBV-negative DLBCL. EBV-positive tumors have fewer features than their EBV-negative counterparts (**: p-val < 0.01, Mann-Whitney U test). B) ISOMAP of DLBCL tumors based on LymphGen subtype-defining features. Unclassified (Other) EBV-positive tumors (X) form a cluster, potentially a novel subtype.

### EBV-positive DLBCL cell line characterization

To better define how well EBV-positive DLBCL cell lines model actual tumors, we conducted RNA-seq, whole exome sequencing (WES), and low-pass whole genome sequencing (WGS) on the 5 published EBV-positive DLBCL lines: Val, Farage, BCKN1, IBL1, and IBL4^34,51–56^. LymphGen mutation features were identified using WES (Table 1, Supplementary Table S7). By RNA-seq, Val exhibited low levels of EBV gene expression in a pattern consistent with Latency I or II (Figure 5A). The latency expression programs of Farage, BCKN1, IBL1, and IBL4 most closely matched Latency III, except that LMP2A transcription was absent or at very lower levels relative to control lymphoblastoid cell lines (LCLs) (Figures 5A-C).

**Figure 5.**
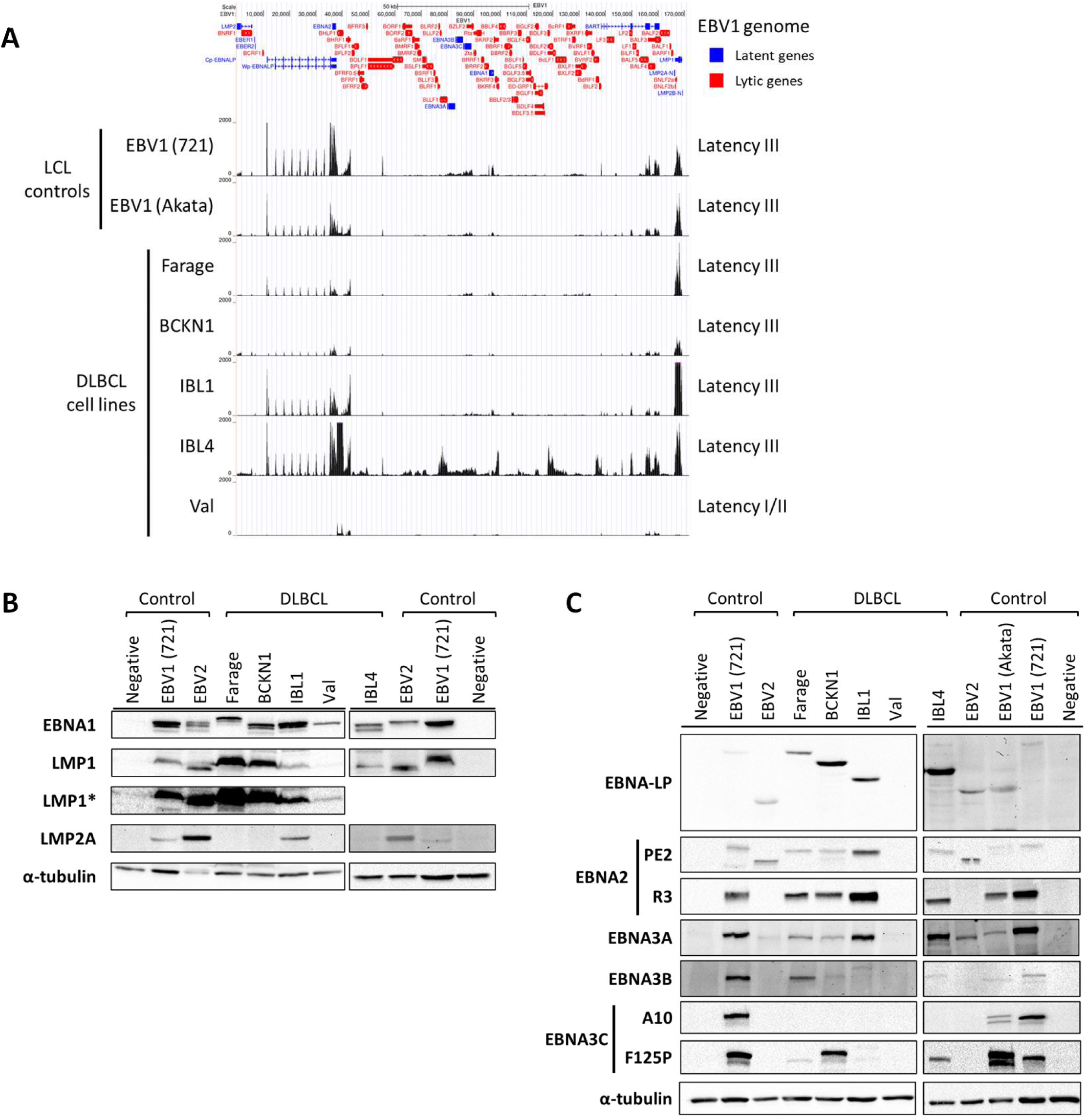
EBV latent gene expression in EBV-positive DLBCL cell lines. A) Tracks displaying RNA-seq data from EBV-positive DLBCL cell lines and control LCLs displayed on the UCSC genome browser. EBV gene annotations for latent (blue) and lytic (red) are shown in the top track. B,C) Immunoblots demonstrating protein expression in EBV-positive DLBCL lines and control LCLs. For some EBV proteins, multiple antibodies were used as indicated. *: longer exposure time.

**Table 1.** LymphGen-defining mutations in EBV-positive DLBCL cell lines.

|  |  | MCD | N1 | A53 | BN2 | ST2 | EZB |
| --- | --- | --- | --- | --- | --- | --- | --- |
| EBV-positive | Farage | CDKN2A,<br>PRDM1, TNRC18 |  | TP53 | PRKCB,<br>TRRAP | DDX3X, PRRC2C,<br>ZNF516 | CREBBP,<br>CIITA,<br>EP300 |
|  | BCKN1 | HLA-A, TNRC18 | IKBKB,<br>NOTCH1 |  |  | ZFP36L1, ITPKB, ACTB |  |
|  | IBL1 |  |  |  |  |  | MAP2K1 |
|  | IBL4 | PIM1 | ALDH18A1,<br>ID3 | TP53 | CCND3 | ACTG1 | EZH2 |

### RNA-seq accurately reflects EBV latent protein expression

To evaluate the accuracy of RNA-seq for latency type, we performed EBV latency protein immunoblotting. We confirmed EBNA1 expression in each EBV-positive DLBCL line (Figure 5B). LMP1 was expressed in Farage, BCKN1, IBL1 and IBL4, but barely detected in Val, despite Val being previously classified as latency II based on LMP1 expression^34^. With the exception of IBL1, LMP2A protein was not detected in the DLBCL lines, but readily detected in LCLs infected with either EBV1 or EBV2 (Figure 5B and Supplementary Figure S4B). Examination of the predicted LMP2A protein sequence from our RNA-seq found that the epitopes for LMP2A monoclonals harbored sequence variations (Supplementary Figure S4A). In reconstructing the LMP2A mRNA from IBL1, we discovered that many spliced aberrantly from LMP2A exon 1A into BBLF2/3 exon 2. We were able to confirm this fusion transcript by RT-PCR (Supplementary Figure S5), but did not observe a corresponding protein product. Despite the sequence variations and splicing aberrancies, it is possible that IBL1 expresses a functional LMP2A protein. Nevertheless, it is clear that LMP2A expression is not a consistent feature of EBV-positive DLBCL cell lines – even those that appear to exhibit a latency III-like pattern.

Latency III-specific nuclear proteins were also assessed by immunoblot (Figure 5C). As expected for a latency I/II cell line, Val was negative for each of these proteins. EBNA-LP and EBNA2 blotting confirmed their expression in each of the Latency III-like DLBCL lines. Blotting for EBNA3 proteins raised several caveats. As expected, Farage, BCKN1, IBL1, and IBL4 were all positive for EBNA3A. However, EBNA3B was only detected strongly in Farage and weakly in BCKN1 and IBL4. EBNA3C was detectable in Farage, BCKN1, and IBL4 only with the polyclonal anti-EBNA3C antibody. This variable detection correlated with epitope-disrupting sequence variations in the EBNA3 proteins (Supplementary Figure S4C). Our results demonstrate that RNA-seq reliably determines latent gene expression patterns and highlights the limitations of relying exclusively on antibodies to define latency types.

### IBL1 is infected with a rare intertypic EBV recombinant

We performed multiple sequence alignments on the predicted protein sequences of the EBNA2 and EBNA3 proteins for each cell line and control LCLs. This revealed that Farage, BCKN1, and Val are infected with EBV1 strains. By contrast, IBL1 had EBNA3A, EBNA3B, and EBNA3C genes from EBV2, but an EBNA2 gene corresponding to EBV1 (Figure 6A-C, Supplementary Figure S6). Western blotting with the EBV1-specific EBNA2 antibody, R3, confirmed that IBL1’s EBNA2 is that of an EBV1 strain (Figure 5C). Thus, IBL1 is infected with an EBV1/EBV2 intertypic recombinant. Such recombinants have on rare occasions been sequenced (<1% of genomes assessed in a study of 241 EBV genomes obtained worldwide^57^), but IBL1 is, to the best of our knowledge, the only cell line harboring such a recombinant. Thus, IBL1 provides a unique opportunity to study the biology of this naturally occurring rare EBV genotype.

**Figure 6.**
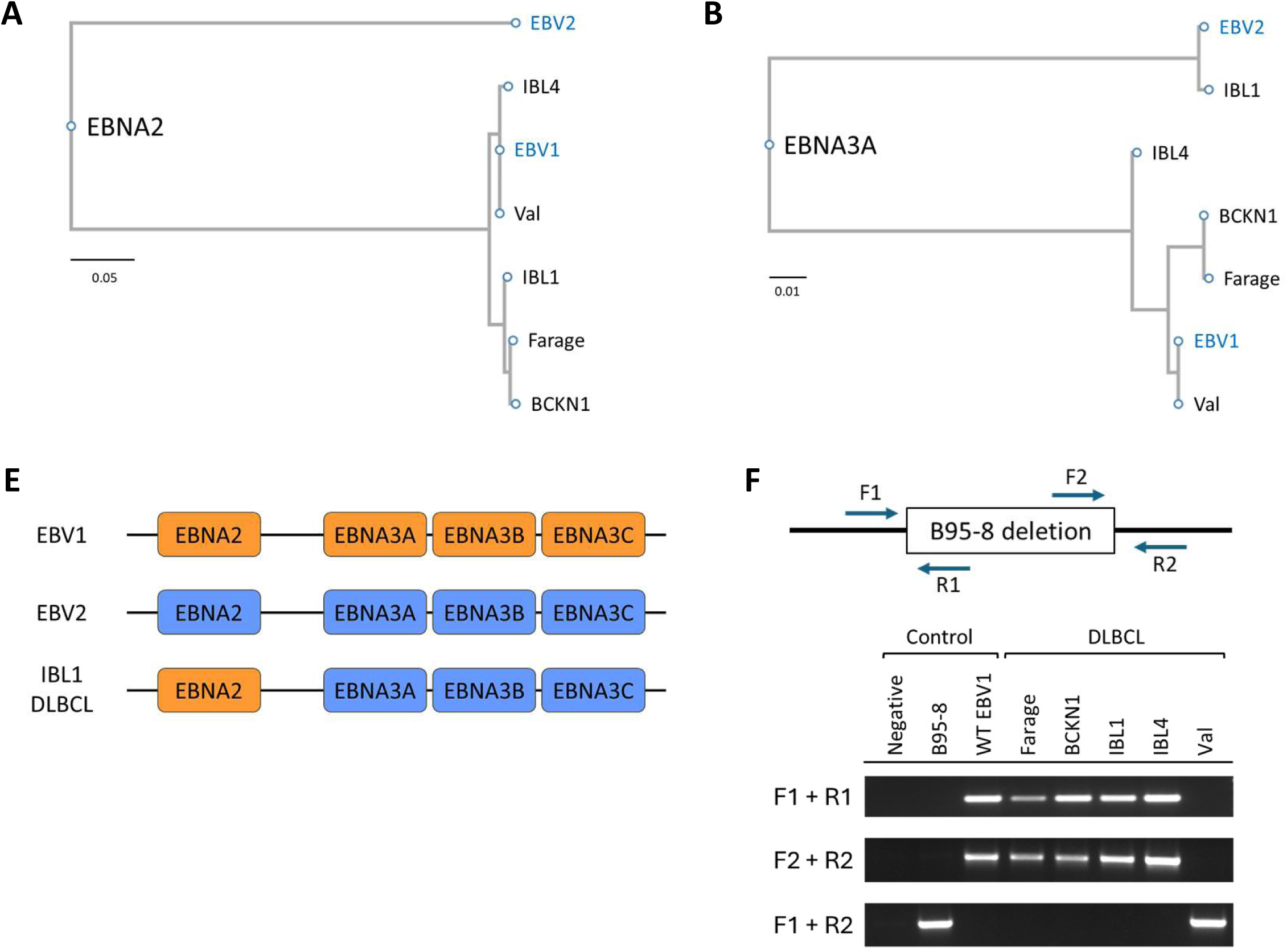
IBL1 is infected with an intertypic EBV1/EBV2 recombinant and Val is not a *bona fide* EBV-positive DLBCL cell line. A, B) Phylogenetic trees for the indicated EBNA proteins are shown based on the predicted protein sequences obtained from *in silico*-translation of sequencing data from the indicated EBV-positive DLBCL lines and EBV1 (NC_007605.1) and EBV2 (NC_009334.1) reference genomes. C) Cartoon representation of the intertypic EBV1/EBV2 recombinant in the IBL1 cell line. D) Diagram showing locations of PCR primers designed to detect the B95-8 deletion region, and representative PCR results (from 3 replicates) using the indicated primer pairs. Controls: BJAB (EBV negative); 721 LCL (B95-8 EBV); Akata LCL (wildtype EBV1).

### Val is not a bona fide EBV-positive DLBCL cell line

An unexpected feature of the RNA-seq data from Val, was the absence of reads mapping to several exons of the BART transcript. Closer inspection showed that the missing sequence corresponded to a ∼12kb deletion specific to the B95-8 laboratory strain of EBV^58^ (Supplementary Figure S7A). Further examination of the LMP2A sequence of Val identified specific tyrosine (Y23) and serine (S444) residues present in B95-8, but absent in all sequenced circulating EBV strains (Supplementary Figure S7B). To confirm this finding, we performed PCR on Val DNA (Figure 6D). Sequencing of this PCR product confirmed that Val harbors a deletion identical to B95-8 (Supplementary Figure S7C). These findings indicate that Val is not an EBV-positive DLBCL, but a cell line infected with the B95-8 EBV laboratory strain *ex vivo*.

## Discussion

Historically, diffuse histiocytic lymphoma (which included DLBCL) was not considered EBV-associated^59^. However, it is now generally accepted that 5-15% of DLBCLs are EBV-positive, and we show that EBV-positive DLBCL, like EBV-negative DLBCL, is not a single disease. An underlying assumption of the LymphGen algorithm is that specific co-occurring mutations collaborate to drive a normal B cell down specific pathways to lymphoma. It is therefore not surprising that EBV oncogenes may substitute for some oncogenic mutations but not others. The observation that many LymphGen subtypes share a mutation spectrum with distinct low-grade lymphomas that can progress to DLBCL predicts that if EBV can cause a specific DLBCL subtype it should also be associated with the corresponding low-grade lymphoma. Thus, one would predict that EBV should be associated with the marginal zone lymphoma (MZL), which shares a mutation spectrum with BN2^4,60^. Indeed, numerous reports have found that EBV is associated with extranodal MZLs, particularly in immunosuppressed persons^61–63^.

Our analysis identified 6 EBV-positive BN2 DLBCL tumors and 11 among the unclassified cases. Some of the latter appear to be in proximity to the BN2 cluster. In such cases, EBV LMP1 substitution for BCL10 or other mutations may have led to “misclassification.” Eight unclassified EBV-positive tumors appear distinct and may represent a novel subtype. Thus, EBV-positive DLBCL likely comprises at least two distinct groups: BN2 and non-BN2. An important question for future study is the extent to which these groups account for the heterogeneity of EBV-positive DLBCL. For example, prior studies have variably reported EBV-positive DLBCL to be associated with either a favorable or unfavorable prognosis compared to EBV-negative disease^7,27–31^. In our analysis, we observed a strong trend toward an unfavorable prognosis within the BN2 EBV-positive subtype, a pattern not seen in the “Other” group (Supplementary Figure S2).

ST2 may also be EBV-associated. Although our analysis only identified one such tumor, this is consistent with expectation given the rarity of ST2, even if the subtype were moderately enriched for EBV-positive cases. Importantly, because LymphGen does not account for the influence of EBV gene products, the detection of any ST2 EBV-positive DLBCL tumors may be important. Activation of JAK/STAT signaling – a hallmark of ST2 – has been repeatedly observed in EBV-positive DLBCL cohorts^8,9,11,64^. In addition, ST2 DLBCL shares a mutation spectrum with nodular lymphocyte predominant Hodgkin lymphoma (NLP-HL), which is EBV-associated, albeit less strongly than classical HL^65,66^. Similarly, T-cell/histiocyte-rich large B cell lymphoma (THR-LBCL) not only shares a mutation spectrum with ST2 but also resembles EBV-positive DLBCL morphologically^28^. While THR-LBCL has not been directly linked to EBV, this is essentially by definition, as EBER positive THR-LBCL cases are classified as EBV-positive DLBCL^67^. Thus, although we did not find ST2 to be enriched for EBV-positive DLBCL, we cannot exclude this possibility, and multiple lines of evidence suggest biological parallels between ST2 and EBV-positive DLBCL that warrant further investigation.

In contrast to other EBV-associated cancers, DLBCL does not exhibit a stereotyped pattern of EBV latent gene expression. Some of this heterogeneity likely reflects that EBV-positive DLBCL is not a single disease entity. However, our results show that even within the BN2 subtype, latency II and latency III patterns occur with equal frequency. Perhaps these tumors expressed latency III initially and, during tumor evolution, acquired cellular mutations that allowed continued proliferation under the less immunogenic latency II program.

Intriguingly, we observed loss-of-function CD70 mutations only in EBV-positive BN2 tumors expressing latency III. Inherited CD70 mutations confer marked susceptibility to EBV-associated lymphomas due to impaired T cell surveillance of EBV-infected B cells^38,39^. By analogy, these somatic CD70 mutations may alleviate immune surveillance pressure that would otherwise force EBV-positive BN2 tumors toward the more restricted latency II program, thereby permitting continued expression of the more transforming latency III program. An important unanswered question is what driver mutations BN2 tumors must acquire to proliferate while expressing the more-restricted latency II. Although mutations in the Notch pathway are an obvious possibility—given that the Notch-interacting EBNA proteins are expressed only in latency III—we did not detect a difference in Notch pathway mutation frequency between latency II and latency III BN2 tumors (Supplementary Table S3B).

An unexpected feature of EBV latent gene expression in DLBCL was the absence of LMP2A expression in tumors and most cell lines. For simplicity, we designated tumors as latency II or latency III if they expressed the other genes associated with those programs, even when LMP2A was absent. Although deviations from canonical latent gene expression are not rare, our observation that LMP1 is expressed more consistently than LMP2A is distinctive. In most EBV-associated tumors, such as NPC, EBV-associated gastric carcinoma (EBVaGC), and even BL, LMP2A is detected in a much higher proportion of tumors than LMP1^68^. Because both membrane proteins act as oncogenes and immune surveillance targets, their expression confers both benefits and risks to an EBV-associated tumor. The mechanisms underlying the preferential loss of LMP2A in DLBCL warrant study.

EBV-positive DLBCL cell lines are important reagents for testing functional interactions between host mutations and viral oncogenes, but it is essential to distinguish them from lines superinfected with EBV in the laboratory, where the virus played no role in lymphomagenesis. Because Val was the only non-latency III cell line, the discovery that it carries the B95-8 laboratory strain is disappointing. In retrospect, the claim that it was “derived from the bone marrow of a 50-year-old woman with B-acute lymphoblastic leukemia (B-ALL) in 1985” seems an improbable origin for a DLBCL line^53,69,70^. Clarifying the provenance of Val and preventing further adulteration of the EBV-positive DLBCL literature is an important contribution of our study, establishing a firmer foundation for understanding EBV’s role in DLBCL.

Because each DLBCL subtype harbors distinct driver mutations, EBV’s contribution may vary by subtype. We were able to validate cell lines that model either the ST2 (Farage) or Other (BCKN1, IBL1, IBL4) subtypes (Table 1, Supplementary Table S7). However, the lack of EBV-positive BN2 cell lines presents a significant obstacle to defining which EBV latency genes drive BN2 proliferation and survival. Establishing additional EBV-positive DLBCL cell lines that more accurately reflect subtypes associated with EBV infection remains an important goal. Some insights may still be gleaned from the study of Riva, an EBV-negative BN2 cell line^4^. For example, our results suggest that LMP1 may supplant the need for NF-κB activating mutations, a hypothesis that could be tested by examining whether stable expression of LMP1 in Riva reduces its dependence on the CARD11-BCL10-MALT1 complex. Ultimately, clarifying these relationships will be critical for understanding how EBV reshapes the genetic requirements of specific DLBCL subtypes.

Our cell line validation yielded additional insights. Post-transcriptional mechanisms influence whether, and to what extent, an mRNA is translated into protein^71,72^. Western blotting not only confirmed the accuracy of RNA-seq for predicting EBV gene expression in DLBCL but also demonstrated the value of using multiple antibodies to assess EBV latency proteins. We encountered several false negatives due to lack of epitope conservation, indicating that RNA-seq is the more robust and less error-prone approach for determining latency programs. A second unexpected finding was that IBL1 is an intertypic recombinant between EBV1 and EBV2. While such recombinants have been documented in large-scale sequencing efforts, they have not previously been isolated in culture. Thus, IBL1 represents a unique resource for studying the biology of this unusual hybrid EBV genome.

Our findings underscore that EBV-positive DLBCL is not a single disease but instead comprises biologically distinct subgroups that cannot be fully captured by classification systems built on EBV-negative tumors. By showing how EBV oncogenes may substitute for, or collaborate with, host mutations in a subtype-specific manner, we highlight the need for classification schemes that explicitly account for EBV’s contribution to lymphomagenesis. This perspective reframes EBV not as a passive passenger but as an active determinant of tumor biology, shaping both the mutational landscape and therapeutic vulnerabilities of DLBCL. Moving forward, the integration of EBV status, viral gene expression, and host genetics will be essential to define the full spectrum of EBV-positive DLBCL and to identify strategies for targeted interventions. In this way, EBV-positive DLBCL can serve as a model for understanding how tumor viruses intersect with cancer genetics to drive disease.

## Materials and Methods

### Data resource

RNA sequencing and whole exome sequencing (WES) BAM files of tumor samples from Schmitz et al.^3^ were downloaded from the GDC Data Portal (project ID: NCICCR-DLBCL). Additional molecular and clinical information were downloaded from dbGaP (study accession: phs001444.v2.p1).

### RNA sequencing analysis of EBV transcriptomes from tumor data

Primary reads mapping in proper pairs to the EBV and human genomes were extracted from BAM files using SAMtools^73^. EBV-to-human read ratios were calculated and plotted in Python (v3.12.7) using JupyterLab. BAM files of candidate EBV-positive tumors were converted into unaligned BAM (UBAM) files and realigned to the GDC reference genome GRCh38.d1.vd1 with the following modification: chrEBV was replaced with a renumbered version of EBV^74^, in which the origin has been reset such that no transcripts span the origin. Realignment was performed using STAR^75^ v2.7.5c with the same parameters as was used on the GDC BAM files. Reads mapping to the renumbered EBV genome were converted into wig tracks with standard EBV genome coordinates, and visualized using the UCSC Genome Browser^76^.

### Statistics

Mstat v7.0 was used for statistical analysis of the association between LymphGen subtypes and EBV status with Barnard’s exact test with F order (N. Drinkwater, McArdle Laboratory for Cancer Research, School of Medicine and Public Health, University of Wisconsin) and is available for download (https://oncology.wisc.edu/mstat/). Mann-Whitney U test was performed with SciPy in Python in JupyterLab.

### Survival analysis

Clinical and LymphGen subtype information were obtained from Wright et al.^4^. Kaplan-Meier analyses were performed in R (https://www.r-project.org/) version 4.3.2 using the survival (https://CRAN.R-project.org/package=survival) and survminer (https://cran.r-project.org/web/packages/survminer/) packages.

### Differential mutation analysis of cellular mutations

Variant calls from Schmitz et al.^3^ were analyzed in R to determine mutation frequencies in EBV-positive and EBV-negative tumors. For mutations depleted in EBV-positive BN2, the following filters were applied: mutation frequency ≥ 0.1 in EBV-negative BN2, and wildtype in EBV-positive BN2. For mutations enriched in EBV-positive BN2, the following filters were applied: mutation frequency ≥ 0.2 in EBV-positive BN2, and wildtype in EBV-negative BN2 or having a ratio 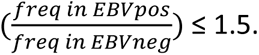 Filtered mutation frequencies were visualized using R. The list of mutated genes were used for pathway enrichment analyses, performed using MSigDB^77^, KEGG Mapper^78^, and Metascape^79^.

### Cell composition analysis using CIBERSORTx

RNA-seq BAM files were quantified using featureCounts, the counts converted into TPM in Python, and the result analyzed using CIBERSORTx^48^ with the Impute Cell Fractions module and the LM22 signature matrix.

### Clustering analysis

ISOMAP was used to visualize the relationship between DLBCL tumors. Processed segment copy number, gene copy number, gene mutational, and translocation datasets were downloaded from dbGaP (phs001444). LymphGen subtype-defining features were extracted to generate an array of binary feature vectors. The Hamming distance between the feature vectors of each tumor in the array was calculated and the epsilon parameter tuned to generate a manifold containing all tumors. The undirected graph was simplified through a shortest-path graph search followed by PCA to generate the visualization.

### Cell lines and culture

All cell lines were maintained in RPMI 1640 Medium (Gibco) supplemented with 10% (BJAB, 721, IBL1, and IBL4) or 20% (Farage, BCKN1, Akata-LCL, and AG876 LCL) fetal bovine serum (FBS) (HyClone Laboratories). All cell culture media were supplemented with 100 U/ml penicillin and 100 μg/ml streptomycin (Gibco). All cells were grown at 37°C in a 5% CO2, humidified atmosphere. Farage, BCKN1, IBL1, IBL4, and Val are EBV-positive DLBCL cell lines and have been previously described^34,51–56^. Farage was obtained from the American Type Culture Collection (ATCC), IBL1, IBL4, and BCKN1 were kindly provided by Ethel Cesarman (Weill Cornell Medical College), Val and additional vials of BCKN1 were kindly provided by Sumita Bhaduri-McIntosh (University of Florida). Each DLBCL cell line was validated using short tandem repeats (STRs) and human leukocyte antigen (HLA) profiles (Supplementary Table 2 & 3). These matched existing profiles for Farage and Val and, for cell lines lacking reference profiles (BCKN1, IBL1, and IBL4), excluded cross-contamination with other known cell lines.

### STR and HLA analyses of cell lines

DNA was isolated from all five EBV-positive DLBCL cell lines using a DNeasy Blood & Tissue Kit (QIAGEN). Short tandem repeat (STR) analyses were performed by the Translational Research Initiatives in Pathology (TRIP) lab at UW-Madison and verified against ATCC and DSMZ references where available. Aligned RNA data from the RNAseq experiment (described above) was analyzed using arcasHLA^80^ (https://github.com/RabadanLab/arcasHLA) for human leukocyte antigen (HLA) typing and verified against TRON Cell Line Portal (TCLP)^81^ (https://github.com/TRON-Bioinformatics/TCLP) and DSMZ references where available. References were found for Farage and Val. There are currently no available references for IBL1, IBL4, and BCKN1. The STR and HLA profiles of all five lines can be found in Supplementary Tables S2 and S3.

### RNA sequencing and whole exome sequencing (WES) of cell lines

For RNA sequencing, live cells were obtained by Ficoll density separation (Ficoll-Paque™ PREMIUM, Cytiva), washed twice in 1X DPBS, pelleted, and resuspended in TRIzol Reagent (Thermo Fisher) at 2×10^6^ cells per 700μl TRizol. RNA isolation was performed using a Direct-zol RNA kits (Zymo Research) according to the manufacturer’s protocol. Library preparation (poly-A), quality control, and paired-end RNA sequencing were performed by the Yale Center for Genome Analysis using NovaSeq. For whole exome sequencing, cells were counted, pelleted, washed with 1X DPBS and repelleted. DNA isolation was performed using a DNeasy Blood & Tissue Kit (QIAGEN). Library preparation, quality control, and WES sequencing were performed by Novogene using NovaSeq. Cell line RNAseq and WES data were deposited in BioProject PRJNA1442779.

### RNA sequencing analysis of cell line samples

RNAseq FASTQ files were assessed for their quality using FastQC (https://qubeshub.org/resources/fastqc). Reads were aligned using STAR to the GDC reference genome with a renumbered EBV genome and wig track generated as detailed above for the tumor data.

### WES analysis of cell line samples

WES FASTQ files were aligned using BWA-MEM^82^ with the GRCh38 GDC reference genome, recalibrated with ApplyBQSR (GATK4^83^), variant-called using Mutect2 (GATK4), and annotated using SnpEff^84^. Further annotations were performed using SnpSift^85^ with the ClinVar, dbSNP, and gnomAD databases. Variants of HIGH or MODERATE impact and those that are not common population SNPs were filtered using SnpSift.

### Immunoblotting

Cells were counted, pelleted, washed with 1X DPBS, repelleted, and resuspended in RIPA lysis buffer (10mM Tris-Cl pH 8.0, 1mM EDTA, 0.5mM EGTA, 1% Triton X-100, 0.1% sodium deoxycholate, 0.1% SDS, 140mM NaCl, 1X protease-inhibitor cocktail [PIC; cOmplete, Mini, EDTA-free protease-inhibitor cocktail; Roche]) at a concentration of 10^6^ cells per 30μl of buffer. The cell suspension was sonicated using a Qsonica Q700 sonicator at 100 mA for at least 1 min and centrifuged at >13,000xg for 30 min, 4°C, and the supernatant was collected for immunoblotting. The collected supernatant was mixed at a 1:1 ratio with 2X SDS sample buffer (0.12 Tris-Cl pH 6.8, 4% SDS, 20% glycerol, 0.3mM bromophenol blue). Samples were boiled at 95°C for 5 minutes (for LMP2A immunoblot, samples were heated at 70°C for 10 minutes). A volume of 18μl (approximately 3×10^5^ cells) per sample per well was loaded onto a homemade 6%, 8%, or 10% SDS-PAGE gel, the gel was run, and the proteins were transferred to a nitrocellulose membrane (Amersham). The membranes were blocked in Tris-buffered saline (TBS) with 0.1% Tween-20 (TBST) containing 5% milk and incubated with the primary antibody overnight at 4°C. Following the primary antibody incubation, the membranes were washed with TBST and incubated with the secondary antibody for 1h at room temperature. All antibody dilutions were prepared in 5% milk solution in TBST. Membranes were washed with TBST, developed using Pierce enhanced chemiluminescence (ECL) Western blotting substrate (Thermo Scientific) according to the manufacturer’s protocol, and visualized using a ChemiDoc (Bio-Rad). Details of primary and secondary antibodies can be found in Supplementary Table S4.

### RT-PCR and sequencing of LMP2A

100ng of total RNA (from BJAB, 721, or IBL-1) isolated with Direct-zol RNA kit (Zymo) was used as input for first strand synthesis with ProtoScript II First Strand cDNA Synthesis Kit (NEB) using Oligo d(T)23 VN primer following manufacturer’s instructions. 1μL of first-strand product was then used in 25μL PCR reaction with Q5 DNA Polymerase (NEB) (98°C 30s, 35 cycles (98°C 10s, 70°C 20s, 72°C 45s), 72°C 2min, 4°C hold), with the following primers: LMP2A_exon1_F2:CCCACCGCCTTATGAGGACC; LMP2A_exon6_R1:TGCAAAATACTGCCACCAGCG; BBLF2/3_R1:GGGTGGAGGGCCGATATCAC (see also Supplementary Table S5). 5μl of the PCR product was run on a 2% TAE agarose gel and visualized using an Azure Imager (Azure Biosystems). PCR products were cleaned up with PCR Purification Kit (QIAGEN) and sent to Plasmidsaurus for long-read sequencing (Premium PCR service).

### Determination of EBV strains

Consensus EBV sequences were assembled from aligned RNA data (described above) using SAMtools consensus with default settings. Coding sequences of EBV’s latent proteins were manually annotated on each consensus sequence with verification using Basic Local Alignment Search Tool (BLAST) (https://blast.ncbi.nlm.nih.gov/Blast.cgi) and SnapGene (v8.1.1) and translated using MEGA11^86^. Multiple sequence alignment (MSA) of the translated sequences was performed in MEGA11 and ClustalW (https://www.genome.jp/tools-bin/clustalw). Phylogenetic analyses were performed using ClustalW, with the PhyML maximum likelihood method. Reference genome accession numbers are: NC_007605.1 (EBV1) and NC_009334.1 (EBV2).

### Validation PCR of the B95-8 deletion region

DNA was isolated from all five EBV-positive DLBCL cell lines using a DNeasy Blood & Tissue Kit (QIAGEN). PCR reactions were performed using the GoTaq G2 Hot Start Green Master Mix (Promega) (95°C 2min, 35 cycles (95°C 30s, 57°C or 58°C 30s, 72°C 1min), 72°C 1min, 4°C hold), with the primer pairs listed in Supplementary Table S6. Samples were run on a 1.5% TAE agarose gel and visualized using a ChemiDoc (Bio-Rad). For junction sequence verification, isolated DNA samples and the sequencing primer (R2) were sent to Functional BioSciences for PCR reaction and Sanger sequencing. The resulting .ab1 files were assessed in FinchTV (v1.3.1) and aligned to the EBV1 genome (NC_007605.1) using SnapGene (v8.1.1).

## Acknowledgement

The authors thank Bill Sugden, Huy Dinh, Joshua Brand, Shannon Kenney, Joseph N. Paulson, Zachary Whitfield, and Elicia Penuel for helpful discussions.

The data/analyses presented in the current publication are based on the use of study data downloaded from the dbGaP web site, under phs001444.v2.p1.

This work was supported by the National Cancer Institute of the National Institutes of Health (NIH; grants PO1 CA022443, U01 CA275247, R01 CA229673). This work was also supported by the University of Wisconsin Carbone Cancer Center Support Grant P30 CA014520.

## Authorship

Q.R., M.H., and E.J. designed and performed research, analyzed and interpreted data, and reviewed and edited the manuscript. Q.R. and E.J. wrote the manuscript. Q.R. and M.H. performed experiments, collected data, and performed statistical analysis.

