## Supplementary document for "Integrating Epstein-Barr virus (EBV) status into diffuse large B cell lymphoma (DLBCL) genetics"

### Supplementary Figure Legends

**Supplementary Figure S1. EBV gene expression tracks of all EBV-positive DLBCL tumors (n=19) identified in the Schmitz et al. cohort.** RNA-seq reads mapping to the EBV genome for each EBV-positive DLBCL tumor are shown, sorted by latency program. EBV latent (blue) and lytic (red) genes are indicated in the annotation (top) track.

**Supplementary Figure S2. Survival analysis of DLBCL cases, considering EBV status.** A) Kaplan-Meier analyses of overall survival (OS) and progression-free survival (PFS) of all (top), BN2 (middle), and Other (bottom) DLBCL with available survival data in the NCI cohort, by EBV status. B) Table summarizing number of DLBCL samples with survival data for each analysis.

**Supplementary Figure S3. Cellular mutation frequency differs with EBV status and latency program.** A) Genes mutated more frequently in EBV-positive BN2. Mutation frequency $\geq$ 0.2 in EBV-positive BN2 and a mutation frequency ratio $(\frac{freq\ in\ EBVpos}{freq\ in\ EBVneg})$ of $\geq$ 1.5. B) Table summarizing the number of DLBCL tumors with mutations in BN2-defining genes (top) and genes mutated more frequently in latency III BN2 tumors (bottom).

**Supplementary Figure S4. Multiple sequence alignments of EBV latent proteins expressed in EBV-positive DLBCL cell lines.** A) LMP2A N-terminal sequences corresponding to the epitopes recognized by the 4E11, 14B7, and 8C3 monoclonal antibodies. Note that LMP2A is polymorphic at residues (63P, 64Y, 79T, 82Q). All EBV+ DLBCL lines exhibited polymorphisms relative to the reference (PYTQ) with BCKN1, IBL1, and IBL4 having PNDP and Farage LDNP[87]. Notably, these polymorphic residues fall partially within the 14B7 epitope. B) Immunoblot for LMP2A protein using the antibodies 14B7 and 4E11. C) EBNA3C sequences corresponding to the epitope recognized by the A10 antibody.

**Supplementary Figure S5. Atypical LMP2A in the IBL1 cell line.** A) Agarose gel showing RT-PCR was performed on RNAs isolated from BJAB (negative control), 721 (positive control), and IBL1 using primer pairs to detect wildtype LMP2A and LMP2A-BBLF2/3 fusion products. Both 721 and IBl1 express wildtype LMP2A, with IBL1 additionally expressing an LMP2A-BBLF2/3 fusion. B) Schematic showing the reconstructed splice junction of the fusion RNA product with exon 1 of LMP2A spliced into exon 2 of BBLF2/3 (i.e. BBLF3), out of frame from the wildtype ORF. C) Immunoblot of LMP2A, LMP1, H3, and α-tubulin from fractionated lysates of a WT EBV1 LCL control (721) and IBL1. Whole-cell lysate (WCL), and the three cellular fractions: cytoplasmic (C), membrane and organelle (M), and cytoskeletal and nuclear (N) are indicated. There was no significant difference in the protein detection of LMP2A in IBL1 and the control LCL and no additional protein signals were detected outside of the WT protein, despite the apparent abundance of the fusion RNA.

**Supplementary Figure S6. Phylogenetic trees for the EBNA3B and EBNA3C proteins of DLBCL cell lines.** As for Figure 6A and 6B, phylogenetic trees were constructed from predicted protein sequences obtained from *in silico*-translation of sequencing data from the indicated EBV-positive DLBCL lines and EBV1 (NC_007605.1) and EBV2 (NC_009334.1) reference genomes.

**Supplementary Figure S7. Additional evidence that Val is not a *bona fide* EBV-positive DLBCL cell line.** A) UCSC genome browser tracks displaying low-pass WGS read coverage of the EBV genome from the indicated EBV-positive DLBCL cell lines. Controls: wildtype EBV1 (Akata LCL), wildtype EBV2 (AG876 LCL), B95-8 (721 LCL). EBV genome annotations indicate latent (blue) and lytic (red) genes. The B95-8 deletion region is annotated in green. Val has a deletion identical to that of B95-8. B) Multiple sequence alignment showing that Val also has two rare LMP2A polymorphisms (Y23 and S444) present in B95-8, but absent in all sequenced circulating EBV strains. 721 LCL is infected with B95-8; EBV1 reference (NC_007605.1) is a pseudo-wildtype sequence primarily based on B95-8, with the deletion region filled in with sequences obtained from the Raji genome. C) Sanger sequencing result demonstrating that the deletion junction in Val is identical to the B95-8 laboratory strain of EBV.

### Supplementary Figure S1

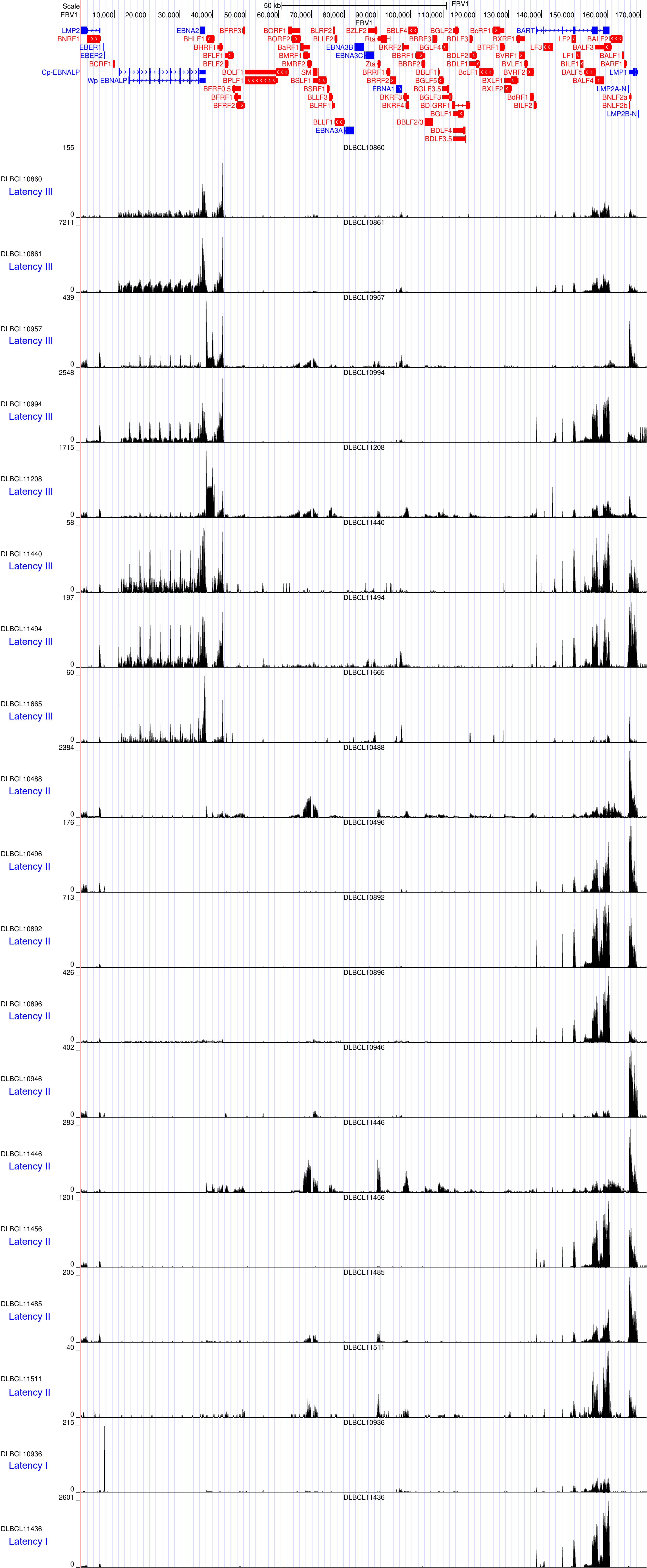
Scale
50 kb
EBV1
DLBCL10860
Latency III
DLBCL10861
Latency III
DLBCL10957
Latency III
DLBCL10994
Latency III
DLBCL11208
Latency III
DLBCL11440
Latency III
DLBCL11494
Latency III
DLBCL11665
Latency III
DLBCL10488
Latency II
DLBCL10496
Latency II
DLBCL10892
Latency II
DLBCL10896
Latency II
DLBCL10946
Latency II
DLBCL11446
Latency II
DLBCL11456
Latency II
DLBCL11485
Latency II
DLBCL11511
Latency II
DLBCL10936
Latency I
DLBCL11436
Latency I

### Supplementary Figure S2

**A**

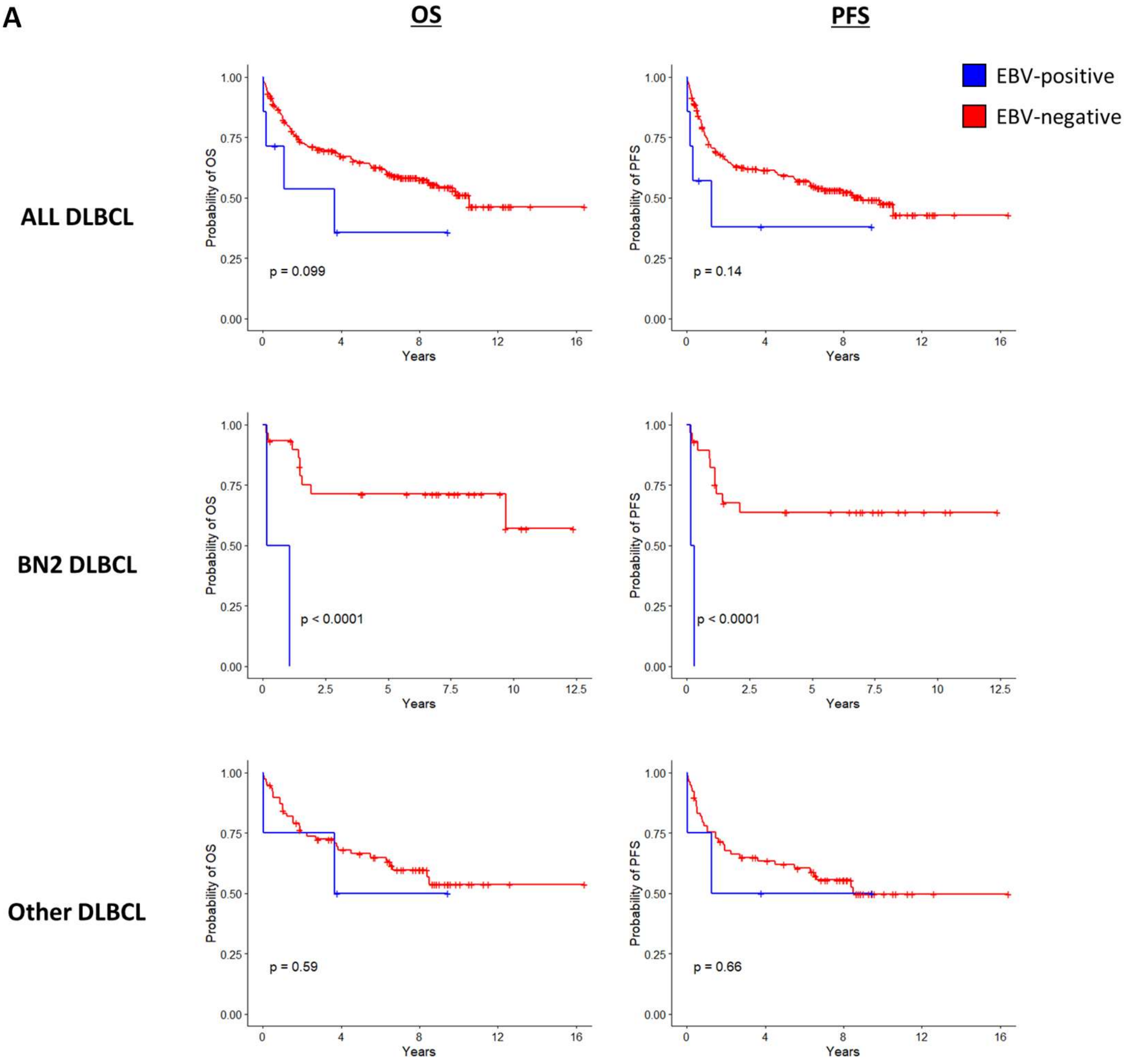

**B**

| | | n for OS | n for PFS |
|---|---|---|---|
| All DLBCL | EBVpos | 7 | 7 |
| | EBVneg | 227 | 222 |
| BN2 | EBVpos | 2 | 2 |
| | EBVneg | 30 | 29 |
| Other | EBVpos | 4 | 4 |
| | EBVneg | 78 | 78 |

### Supplementary Figure S3

**A**

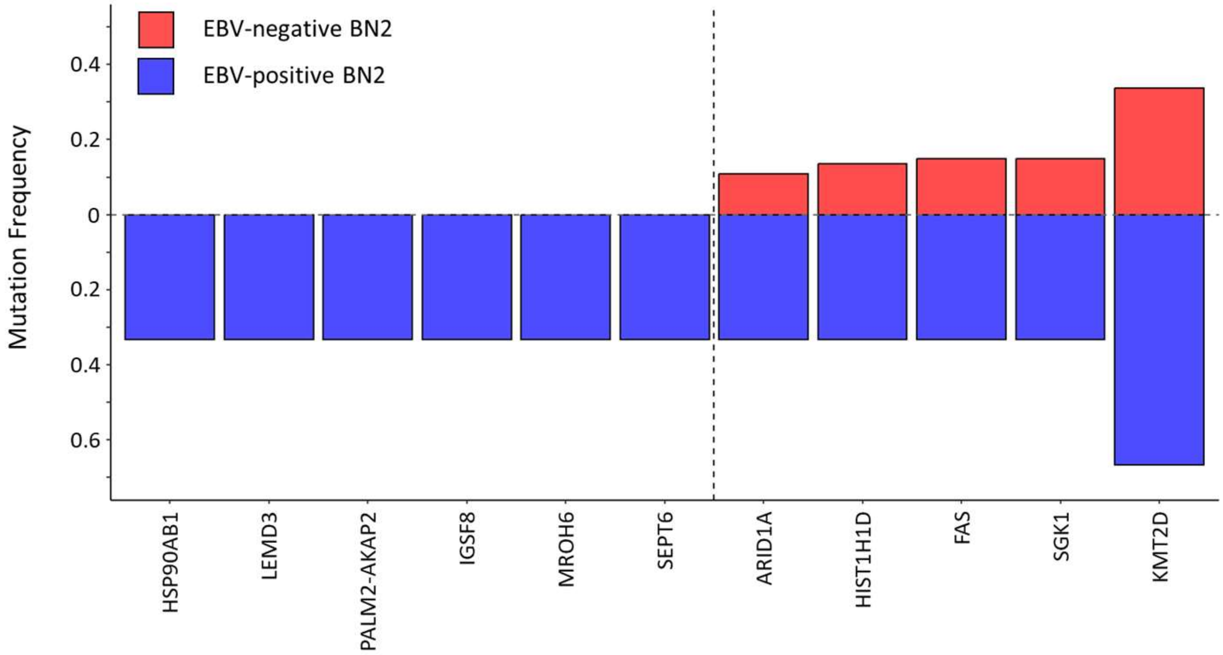

**B**

| | Gene | Tumor count | |
|---|---|---|---|
| | | **Latency 2 (n=3)** | **Latency 3 (n=3)** |
| BN2 Notch mutations | **Notch2** | 2 | 1 |
| | **SPEN** | 1 | 1 |
| | **DTX1** | 1 | 3 |
| Mutations enriched in Latency 3 | **CD70** | 0 | 3 |
| | **CREBBP** | 0 | 2 |
| | **HLA-B** | 0 | 2 |
| | **OSBPL10** | 0 | 2 |
| | **PFKM** | 0 | 2 |

### Supplementary Figure S4

#### A LMP2A antibody epitope

**4E11 epitope**

```
              1                  20                   40
EBV1     MGSLEMVPMGAGPPSPGGDPDGYDGGNNSQYPSASGSSGN
EBV2     MGSLEMVPMGAGPPSPGGDPDGDDGGNNSQYPSASGSSGN
Farage   MGSLEMVPMGAGPPSPGGDPDGDDGGNNSQYPSASGSSGN
BCKN1    MGSLEMVPMGAGPPSPGGDPDGDDGGNNSQYPSASGSSGN
IBL1     MGSLEMMPMGAGPPSPGGDPDGDDGGNNSQYPSASGSSGN
IBL4     MGSLEMVPMGAGPPNPGGDPDGDDGGNNSQHPSVSGSPGN
Val      MGSLEMVPMGAGPPSPGGDPDGYDGGNNSQYPSASGSSGN
```

**14B7 epitope**

```
              31                  51                   71
EBV1     YPSASGSSGNTPTPPNDEERESNEEPPPPYEDPYWGNGDRH
EBV2     YPSASGSSGNTPTPPNDEERESNEEPPPPYEDPYWGNGDRH
Farage   YPSASGSSGNTPTPPNDEERESNEEPPPPYEDLDWGNGDRH
BCKN1    YPSASGSSGNTPTPPNDEERESNEEPPPPYEDPDWGNGDRH
IBL1     YPSASGSSGNTPTPPNDEERESNEEPPPPYEDPDWGNGDRH
IBL4     HPSVSGSPGNTPTPPNDEERESNEEPPPPYEDPDWGNGDRH
Val      YPSASGSSGNTPTPPNDEERESNEEPPPPYEDPYWGNGDRH
```

**8C3 epitope**

```
              75                  95                   115
EBV1     QPLGTQDQSLYLGLQHDGNDGLPPPPYSPRDDSSQHIYEEA
EBV2     QPLGTQDQSLYLGLQHDGNDGLPPPPYSPRDDSSQHIYEEA
Farage   QPLGNQDPSLYLGLQHDGNDGLPPPPYSPRDDSSQHIYEEA
BCKN1    QPLGNQDPSLYLGLQHDGNDGLPPPPYSPRDDSSQHIYEEA
IBL1     QPLGNQDPSLYLGLQHDGNDGLPPPPYSPRDDSSQHIYEEA
IBL4     QPLGNQDPSLYLGLQHDGNDGLPPPPYSPRDDSSQHIYEEA
Val      QPLGTQDQSLYLGLQHDGNDGLPPPPYSPRDDSSQHIYEEA
```

## B

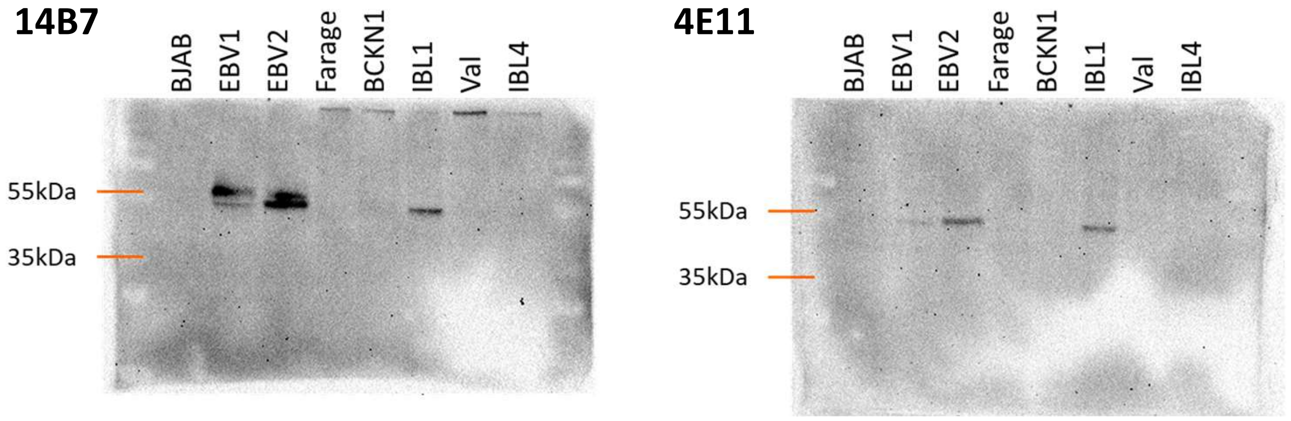

#### C EBNA3C antibody epitope

**A10 epitope**

```
              661                 680                  700
EBV1     MQQEPSSHLQSATQPTTPRPSWAPSVCALSVMDAGKAQPI
EBV2     IQQEPSSQQQPATQSTPPCQSWVPSVYVLPAVDAGNAQPL
Farage   MQQEPSSHLQSATQPTTPRPSWVPSVCALSVMDAGKAQPI
BCKN1    MQQEPSSHLQSATQPTTPRPSWVPSVCALSVMDAGKAQPI
IBL1     IQQEPSSQQQPATQYTPPCQSWVPSVYVLPAVDAGNAQPL
IBL4     MQQEPSSHLQSATQPTMPRPSWVPSVCALSVMDAGKAQPI
Val      MQQEPSSHLQSATQPTTPRPSWAPSVCALSVMDAGKAQPI
```

### Supplementary Figure S5

**A**

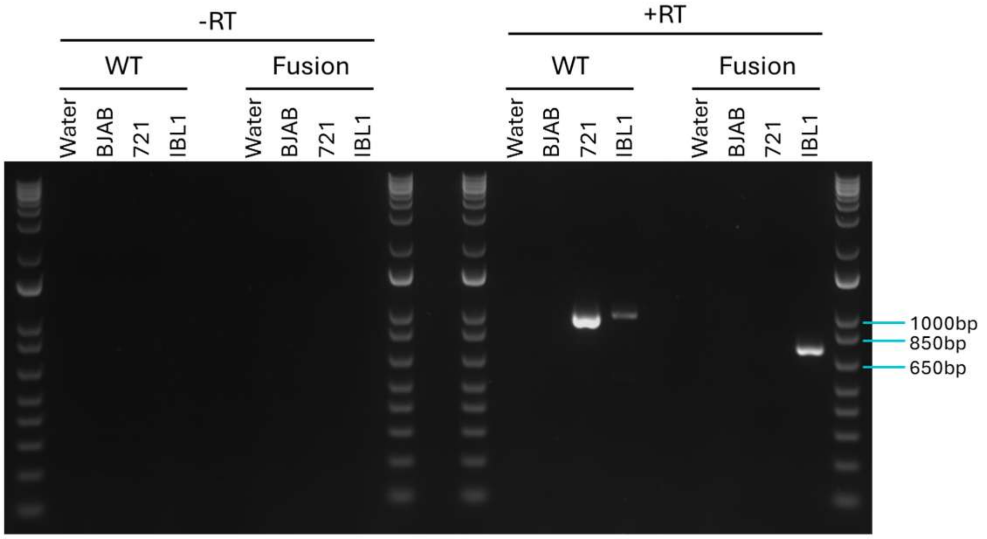

WT : LMP2A wildtype
Fusion : LMP2A-BBLF/23 fusion

**B**

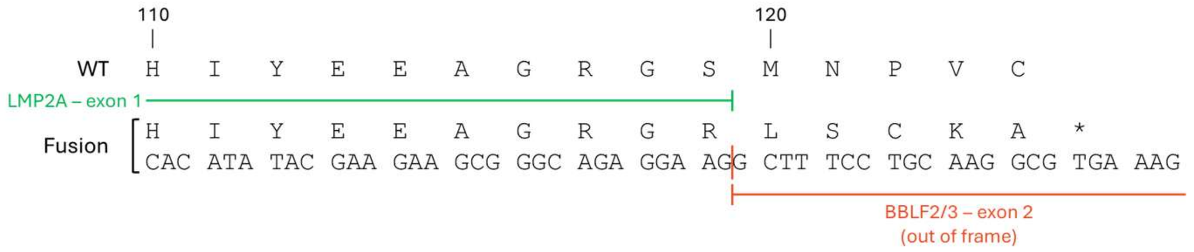

**C**

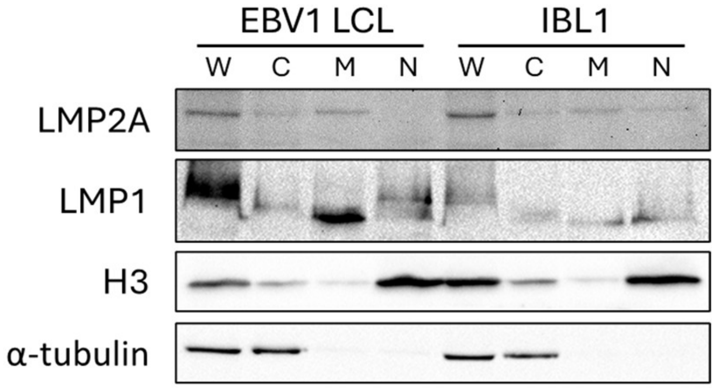

### Supplementary Figure S6

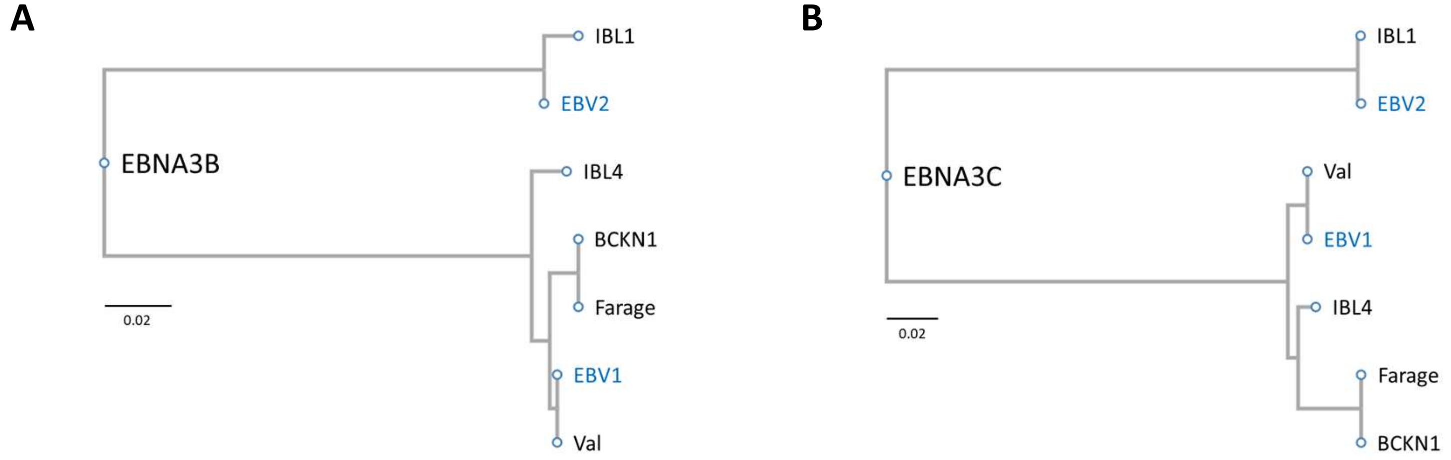

### Supplementary Figure S7

**A**

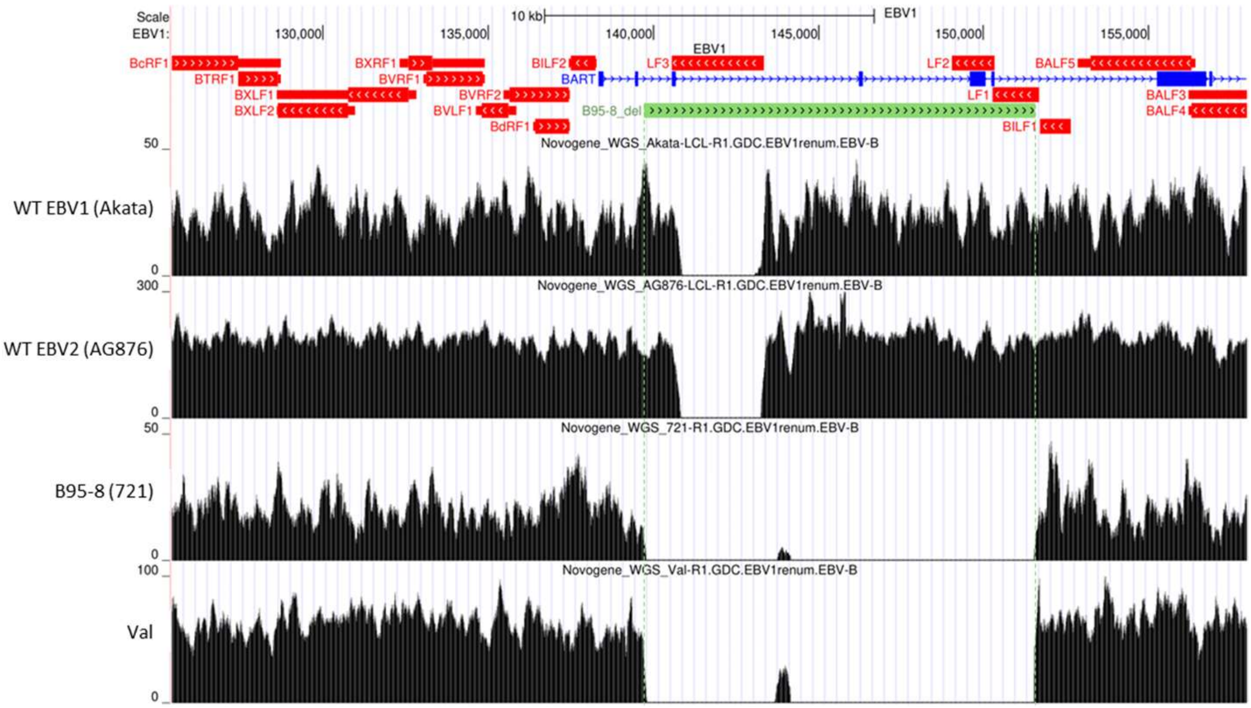

**B**

```
                1                   20  23            40
B95-8           MGSLEMVPMGAGPPSPGGDPDGYDGGNNSQYPSASGSSGN
EBV1 (Akata)    MGSLEMVPMGAGPPSPGGDPDGDDGGNNSQYPSASGSSGN
EBV2            MGSLEMVPMGAGPPSPGGDPDGDDGGNNSQYPSASGSSGN
Farage          MGSLEMVPMGAGPPSPGGDPDGDDGGNNSQYPSASGSSGN
BCKN1           MGSLEMVPMGAGPPSPGGDPDGDDGGNNSQYPSASGSSGN
IBL1            MGSLEMMPMGAGPPSPGGDPDGDDGGNNSQYPSASGSSGN
IBL4            MGSLEMVPMGAGPPNPGGDPDGDDGGNNSQHPSVSGSPGN
Val             MGSLEMVPMGAGPPSPGGDPDGYDGGNNSQYPSASGSSGN

                411                 430           444 450
B95-8           TEWGSGNRTYGPVFMCLGGLLTMVAGAVWLTVMSNTLLSA
EBV1 (Akata)    TEWGSGNRTYGPVFMCLGGLLTMVAGAVWLTVMTNTLLSA
EBV2            TEWGSGNRTYGPVFMCLGGLLTMVAGAVWLTVMTNTLLSA
Farage          TEWGSGNRTYGPVFMCLGGLLTMVAGAVWLTMMTNTRLSA
BCKN1           TEWGSGNRTYGPVFMCLGGLLTMVAGAVWLTVMTNTLLSA
IBL1            TEWGSGNRTYGPVFMCLGGLLTMVAGAVWLTVMTNTLLSA
IBL4            TEWGSGNRTYGPVFMCLGGLLTMVAGAVWLTVMTNTLLSA
Val             TEWGSGNRTYGPVFMCLGGLLTMVAGAVWLTVMSNTLLSA
```

**C**

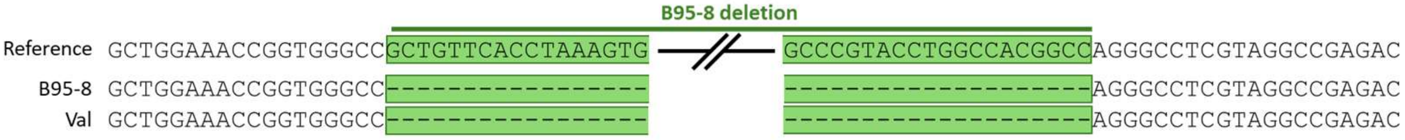

### Supplementary Tables

#### Supplementary Table S1

EBV latent gene expression patterns in EBV-positive tumors from the Schmitz et al. cohort, by LymphGen subtype.

| | | LymphGen subtype | | | | |
|---|---|---|---|---|---|---|
| | | BN2 | MCD | ST2 | Other | Total |
| EBV latency | III | 3 | 1 | 1 | 3 | 8 |
| | II | 3 | 0 | 0 | 6 | 9 |
| | I | 0 | 0 | 0 | 2 | 2 |
| | Total | 6 | 1 | 1 | 11 | 19 |

**Supplementary Table S2. Short tandem repeat (STR) profiles of EBV-positive DLBCL cell lines**

| | Farage | BCKN1 | IBL1 | IBL4 | Val |
|---|---|---|---|---|---|
| **FGA** | 20 ,23 | 23,25 | 24,24 | 25,25 | 22,24 |
| **TPOX** | 9,9 | 8,8 | 8,11 | 8,11 | 8,8 |
| **D8S1179** | 12,12 | 13,13 | 13,14 | 13,13 | 11,12 |
| **vWA** | 14,15 | 17,19 | 17,19 | 18,18 | 14,15 |
| **Amelogenin** | X,X | X,Y | X,Y | X,X | X,X |
| **Penta_D** | 9,12 | 13,13 | 8,10 | 12,13 | 9,9 |
| **CSF1PO** | 11,12 | 11,11 | 12,12 | 10,11 | 10,12 |
| **D16S539** | 11,12 | 13,13 | 12,12 | 11,13 | 9,13 |
| **D7S820** | 12,12 | 8,10 | 12,13 | 10,10 | 10,10 |
| **D13S317** | 11,13 | 12,12 | 11,11 | 11,13 | 10,14 |
| **D5S818** | 12,12 | 12,12 | 11,12 | 8,13 | 11,11 |
| **Penta_E** | 7,18 | 11,15 | 14,22 | 8,11 | 7,7 |
| **D18S51** | 12,13 | 14,14 | 15,18 | 12,17 | 16,17 |
| **D21S11** | 29,29 | 30,32.2 | 29,29 | 30,32 | 27,30 |
| **TH01** | 8,9 | 6,9.3 | 9,9.3 | 9,9 | 6,9 |
| **D3S1358** | 14,18 | 18,18 | 17,18 | 15,15 | 17,19 |
| **Matches** | 100% match<br>Farage (ATCC CRL-2630) | N/A | N/A | N/A | 100% match<br>Val (DSMZ ACC-586) |

**Supplementary Table S3. Human leukocyte antigen (HLA) profiles of EBV-positive DLBCL cell lines** HLA p-groups reported below have been validated in at least two replicates.

| | Farage | BCKN1 | IBL1 | IBL4 | Val |
|---|---|---|---|---|---|
| **A1** | A*03:01P | A*03:01P | A*30:02P | A*02:01P | A*02:01P |
| **A2** | A*24:02P | A*03:01P | A*02:05P | A*26:01P | A*68:01P |
| **B1** | B*14:02P | B*35:03P | B*18:01P | B*45:01P | B*15:01P |
| **B2** | B*51:01P | B*35:03P | B*37:01P | B*45:01P | B*15:01P |
| **C1** | C*08:02P | C*04:01P | C*05:01P | C*07:01P | C*03:03P |
| **C2** | C*16:02P | C*04:01P | C*06:02P | C*07:01P | C*03:03P |
| **DMA1** | DMA*01:01P | DMA*01:01P | DMA*01:01P | DMA*01:01P | DMA*01:01P |
| **DMA2** | DMA*01:01P | DMA*01:01P | DMA*01:01P | DMA*01:01P | DMA*01:01P |
| **DMB1** | DMB*01:01P | DMB*01:01P | DMB*01:01P | DMB*01:01P | DMB*01:01P |
| **DMB2** | DMB*01:01P | DMB*01:01P | DMB*01:01P | DMB*01:01P | DMB*01:01P |
| **DOB1** | DOB*01:01P | DOB*01:01P | DOB*01:01P | DOB*01:01P | DOB*01:01P |
| **DOB2** | DOB*01:01P | DOB*01:01P | DOB*01:01P | DOB*01:01P | DOB*01:01P |
| **DPA11** | DPA1*01:03P | DPA1*01:03P | DPA1*02:01P | DPA1*02:02P | DPA1*02:02P |
| **DPA12** | DPA1*01:03P | DPA1*02:02P | DPA1*01:03P | DPA1*01:03P | DPA1*02:02P |
| **DPB11** | DPB1*04:01P | DPB1*02:01P | DPB1*17:01P | DPB1*01:01P | DPB1*19:01P |
| **DPB12** | DPB1*04:01P | DPB1*04:01P | DPB1*02:02P | DPB1*04:01P | DPB1*19:01P |
| **E1** | E*01:01P | E*01:03P | E*01:03P | E*01:01P | E*01:01P |
| **E2** | E*01:03P | E*01:03P | E*01:03P | E*01:01P | E*01:01P |
| **F1** | F*01:01P | F*01:01P | F*01:01P | F*01:01P | |
| **F2** | F*01:01P | F*01:01P | F*01:01P | F*01:01P | |
| **L1** | | | | | |
| **L2** | | | | | |
| **DOA1** | | DOA*01:02P | DOA*01:01P | DOA*01:01P | DOA*01:02P |
| **DOA2** | | DOA*01:01P | DOA*01:01P | DOA*01:01P | DOA*01:02P |
| **DQA11** | | DQA1*03:01P | DQA1*05:01P | DQA1*02:01P | DQA1*01:03P |
| **DQA12** | | DQA1*03:01P | DQA1*03:01P | DQA1*01:01P | DQA1*01:03P |
| **DQB11** | | DQB1*03:02P | DQB1*02:01P | DQB1*02:01P | DQB1*06:03P |
| **DQB12** | | DQB1*03:02P | DQB1*04:02P | DQB1*05:01P | DQB1*06:11P |
| **DRA1** | | DRA*01:01P | DRA*01:01P | DRA*01:01P | DRA*01:01P |
| **DRA2** | | DRA*01:01P | DRA*01:01P | DRA*01:01P | DRA*01:01P |
| **DRB11** | | DRB1*04:05P | DRB1*04:06P | DRB1*07:01P | DRB1*13:01P |
| **DRB12** | | DRB1*04:05P | DRB1*03:01P | DRB1*01:02P | DRB1*13:01P |
| **DRB31** | | | DRB3*02:02P | | DRB3*02:02P |
| **DRB32** | | | DRB3*02:02P | | DRB3*02:02P |
| **K1** | | K*01:01:01 | K*01:01:01 | | |
| **K2** | | K*01:01:01 | K*01:02 | | |
| **J1** | | | | | |
| **J2** | | | | | |
| **Matches** | Farage (ATCC CRL-2630) via Cellosaurus | N/A | N/A | N/A | Val (DSMZ ACC-586) |

#### Supplementary Table S4

**Primary antibodies**

| Protein target | Antibody | Dilution | Host |
|---|---|---|---|
| EBNA1 | IH4 (ascites) | 1:100 | Rat |
| EBNA2 | Abcam PE2 | 1:2500 | Mouse |
| | R3 | 1:5000 | Rat |
| EBNA3A | Exalpha F115P | 1:1000 | Sheep |
| EBNA3B | Exalpha F120P | 1:1000 | Sheep |
| EBNA3C | Exalpha F125P | 1:1000 | Sheep |
| | A10 | 1:3000 | Mouse |
| EBNA-LP | 4D3 | 1:100,000 | Mouse |
| LMP1 | Abcam CS1-4 | 1:200 | Mouse |
| LMP2A | 4E11 | 1:100 | Rat |
| | 14B7 | 1:200 | Rat |
| | 8C3 | 1:100 | Rat |
| α-tubulin | Novus Biologicals NB120-11304 | 1:2000 | Mouse |

IH4 and R3 are gifts from Bill Sugden. 4D3 is a gift from Shannon Kenney. 4E11 and 8C3 is a gift from Richard Longnecker.

**Secondary antibodies**

| Antibody | Source | Dilution |
|---|---|---|
| α-rat HRP | Thermo 31474 or Invitrogen A10549 | 1:2000 |
| α-mouse HRP | Lifetech A24524 | 1:2000 |
| α-sheep HRP | Thermo 31480 | 1:2000 |
| α-rabbit HRP | Invitrogen A10547 | 1:2000 |

#### Supplementary Table S5

**Primers for LMP2A RT-PCR**

| Name | Binding Region | Sequence |
| --- | --- | --- |
| LMP2A_exon1_F2 | LMP2A exon 1 | CCCACCGCCTTATGAGGACC |
| LMP2A_exon6_R1 | LMP2A exon 6 | TGCAAAATACTGCCACCAGCG |
| BBLF2/3_R1 | BBLF2/3 exon 2 | GGGTGGAGGGCCGATATCAC |

**PCR primer pairs**

| | Primer | Expected Product Size (bp) |
| --- | --- | --- |
| WT | LMP2A_exon1_F2 | 963 |
| | LMP2A_exon6_R1 | |
| Fusion | LMP2A_exon1_F2 | 725 |
| | BBLF2/3_R1 | |

#### Supplementary Table S6

**Primers for B95-8 deletion PCR and sequencing**

| Primer Abbreviation | Primer name | Binding region | Sequence |
|---|---|---|---|
| F1 | B95.8_del_F1 | Outside B95-8 deletion | GCCCATCGACGTATCGCTGG |
| F2 | B95.8_del_F3 | Inside B95-8 deletion | AGCGTGCACCTGGAAGATGCAGC |
| R1 | B95.8_del_R3 | Inside B95-8 deletion | TACCTAGGCCTGCGTCCCAC |
| R2* | B95.8_del_R2 | Outside B95-8 deletion | GGAAGCCGCGCCTGATATGC |

*This primer was also used for Sanger sequencing

**PCR primer pairs**

| | Primer | Annealing temperature used in PCR | Expected product size (bp) | |
|---|---|---|---|---|
| | | | With B95-8 deletion | Without B95-8 deletion |
| 1 | F1<br>R1 | 58°C | NA | 303 |
| 2 | F2<br>R2 | 57°C | NA | 568 |
| 3 | F1<br>R2 | 57°C | 465 | (12,296) |

#### Supplementary Table S7

| Cell Line | GENE | CHROM | POS | TYPE | AA_CHG | CLASS | SUBTYPE |
|---|---|---|---|---|---|---|---|
| Farage | TP53 | chr17 | 7674221 | missense | R89W | mut | A53 |
| | TRRAP | chr7 | 99011184 | missense | A3691S | mut | BN2 |
| | PRKCB | chr16 | 24185553 | missense | A570T | mut | BN2 |
| | CREBBP | chr16 | 3767733 | missense | Q940H | mut | EZB |
| | CIITA | chr16 | 10907870 | missense | L892P | mut | EZB |
| | EP300 | chr22 | 41170520 | frameshift | M1470fs | mut | EZB |
| | PRDM1 | chr6 | 106105841 | missense | T575A | mut | MCD |
| | TNRC18 | chr7 | 5388776 | frameshift | A350fs | mut | MCD |
| | CDKN2A | chr9 | 21971054 | missense | A51V | mut | MCD |
| | PRRC2C | chr1 | 171541554 | missense | R1363H | mut | ST2 |
| | ZNF516 | chr18 | 76442001 | missense | A352T | mut | ST2 |
| | DDX3X | chrX | 41343343 | missense | A38V | mut | ST2 |
| BCKN1 | HLA-A | chr6 | 29944325 | frameshift | G276fs | mut | MCD |
| | TNRC18 | chr7 | 5313461 | missense | A2488D | mut | MCD |
| | IKBKB | chr8 | 42322135 | missense | V607A | mut | N1 |
| | NOTCH1 | chr9 | 136502399 | missense | G1753R | mut | N1 |
| | ITPKB | chr1 | 226634750 | missense | R921L | mut | ST2 |
| | ACTB | chr7 | 5527845 | missense | S344F | mut | ST2 |
| | ZFP36L1 | chr14 | 68790015 | missense | A248S | mut | ST2 |
| IBL1 | MAP2K1 | chr15 | 66444668 | missense | L177M | mut | EZB |
| IBL4 | TP53 | chr17 | 7675094 | missense | V41A | mut | A53 |
| | CCND3 | chr6 | 41935950 | missense | I94K | mut | BN2 |
| | EZH2 | chr7 | 148836988 | missense | A19V | mut | EZB |
| | PIM1 | chr6 | 37171335 | missense | V242M | mut | MCD |
| | ID3 | chr1 | 23559261 | missense | P56S | mut | N1 |
| | ID3 | chr1 | 23559360 | missense | A23T | mut | N1 |
| | ALDH18A1 | chr10 | 95616506 | missense | I526V | mut | N1 |
| | ACTG1 | chr17 | 81512269 | missense | A29V | mut | ST2 |
